# Characterizing and comparing missense variants in monogenic disease and in cancer

**DOI:** 10.1101/534693

**Authors:** Yizhou Yin, John Moult

## Abstract

In both monogenic disease and cancer a large fraction of causative mutations are missense, even though these are very different types of disease. Here we examine and compare a number of properties of these mutations in the two classes of disease to determine the extent to which the properties of the mutations are similar or different. Analysis of cancer mutations is complicated by the problem of distinguishing between drivers and passengers. After controlling for this factor in three different ways, we find the following: (1) A very high and similar fraction (~90%) of causal mutations in both diseases are at positions under strong selection pressure. (2) Mutations in structurally disordered regions play a minor role in monogenic disease (only about 10% of mutations are in those regions), but a larger role (about 25%) in cancer, largely because of the higher fraction of disordered regions in cancer driver proteins. (3) A large (~75%) and similar fraction of causal mutations in protein cores in both diseases act by significantly destabilizing three-dimensional structure, implying a large impact on protein function. (4) Cancer oncogene mutations tend to be on the protein surface whereas tumor suppressor and monogenic disease mutations are more common in the core. (5) A surprisingly high fraction (~50%) of mutations in cancer passenger genes are at positions under strong selection pressure.

## Introduction

### 1. Overview

The large amount of genomic data now available for monogenic disease and cancer has vastly expanded our knowledge of which mutations are involved in these diseases [1,2]. 5,291 monogenic diseases involving 3,645 genes are included in the Online Mendelian Inheritance in Man (OMIM) database (Online Mendelian Inheritance in Man, OMIM®. McKusick-Nathans Institute of Genetic Medicine, Johns Hopkins University (Baltimore, MD), 2019/01/21. World Wide Web URL: http://omim.org/). The Human Gene Mutation Database (HGMD) [3] currently contains 193,904 monogenic disease mutations found in 6770 genes. Another widely used database, ClinVar (https://www.ncbi.nlm.nih.gov/clinvar/, 12/2018 [4]), contains 88,850 pathogenic or likely pathogenic mutations in 4,486 genes. Sequencing of many cancer sample exomes and a growing number of complete cancer genomes has revealed the mutation landscape for dozens of cancer types [2,5].

A variety of mutation types may be causative of monogenic diseases or be cancer drivers, including single nucleotide variants (SNVs) resulting in effects on expression and splicing, amino acid substitutions (missense), and premature stop codons; small insertions and deletions (INDELS) [6,7]; and particularly in cancer [6], copy number variants (CNVs, large insertions or deletions, deleting or duplicating one or more genes). Larger scale chromosomal changes also play a role in cancer, where genome instability is common [8]. In monogenic disease, and in contrast to complex trait disease [9–11], very few mutations affecting expression have been identified [4]. Data for non-coding contributions in cancer are only now becoming available. Some clear examples have been identified [12,13], but a clear picture has yet to emerge. In monogenic disease, the majority of known mutations are missense [14], a single nucleotide variants causing an amino acid substitution. In cancer, missense mutations also play a major role [6].

### 2. Missense mutations

In this paper we use computational methods to analyze and compare the role of missense mutations in monogenic disease and cancer. There are three primary motivations. First, as noted above, this class of mutation is the most in common in both types of disease, so that a thorough understanding of its role is worthwhile. Second, unlike most INDELs and copy number variants which have a major impact on protein function and hence disease phenotype, missense mutations range from no effect on protein function to complete loss of activity. The wide range of possible molecular impact presents problems for clinical interpretation. As a result, at present, evidence from computational analysis is given low weight in clinical diagnosis in monogenic disease [15]. Greater understanding of how these mutations influence disease phenotype will help improve the usefulness of the computational methods. Third, now that many instances of these mutations now known in both types of disease, it is possible to perform statistical analyses that provide insight into the molecular mechanisms involved.

### 3. Methods for interpretation of missense mutations

Methods for imputing the disease relevance of missense mutations fall into two classes: Those that rely on the pattern of observed amino acid substitutions at a mutation position both across species and paralogs and as common variants within the human population, and those that make use of structural and other molecular function information. Sequence-based methods usually utilize machine learning, and typical features are related to sequence conservation and the pattern of substitutions at the position of interest [16]. An advantage of these methods is that, provided there is a deep enough, diverse enough, and stable alignment, any mutation can be analyzed (currently 92% of a reference set of HGMD monogenic disease mutations [14] using SNPs3D profile [17]). Further, subject to the assumption below, they are effective for all types of underlying mechanisms, including gain of function (highly relevant for mutations in oncogenes). The disadvantage of sequence-based methods is that they provide no insight into the mechanism by which a mutation is involved in disease. The methods implicitly assume that if a mutation plays a causative role in disease, it will affect Darwinian fitness, and thus tend to be selected against. Since many monogenic diseases are early onset and severe enough to affect reproductive success, that is often a reasonable assumption. Relevance to cancer, where driver mutations promote cell growth in many ways, is less obvious. Although the assumption of a relationship between an effect on fitness and disease phenotype is embedded in the methods, there is usually no formal theoretical framework for calculating fitness impact. Rather machine learning [17–21] or other parameterization [22,23] is used to calibrate the relationship between amino acid substitution patterns and disease phenotypes in an *ad hoc* way. Here we use three well calibrated methods, SNP3D profile [17], Polyphen-2 [20] and CADD [19], to ensure the results are not sensitive to a single approach.

Protein structure and function provide a complementary, more mechanism-oriented approach to identifying disease-relevant mutations. Previous studies have shown that a high fraction of monogenic disease and to some extent cancer tumor suppressors mutations destabilize three dimensional-structure [24–26]: for a reference set of monogenic disease proteins [14], SNPs3D stability [25] assigns 72% as destabilizing, and for a reference cancer set [6], 50%~60% [26]. Thus, methods of estimating the change in free energy difference between the folded and unfolded states introduced by an amino acid substitution play an important role.

In this paper we use SNPs3D_Stability [25] to examine the role of destabilization in both monogenic disease and cancer. The method uses empirical potential terms representing van der Waals interactions, electrostatics, conformational strain, solvent accessibility and local flexibility in a non-linear support vector machine model and was trained using monogenic disease data [14] together with interspecies variants as controls. It has been benchmarked against experimental ΔΔG data and monogenic disease mutations [25]. The method returns a binary yes/no estimate of whether a missense variant destabilizes a structure sufficiently to contribute to monogenic disease, together with score related to the confidence of the assignment.

### 4. Identifying driver mutations

There are well-established databases of causative mutations for monogenic disease [4,14], and although these sources are not error-free [27], they are sufficiently accurate for many statistical purposes. Reliable identification of cancer driver mutations remains a major problem, because of the high background of passenger mutations. The mutation load varies by more than two orders of magnitude among individual samples as well as across cancer types [2,5]. For example, acute myeloid leukemia and some pediatric cancers may carry less than 10 nonsynonymous somatic mutations per tumor, while exogenous mutagen induced cancers such as lung cancer and melanoma typically have an average of around two hundred [5,28,29]. It has been generally accepted that only a small number of the somatic mutations [30,31] (the ‘drivers’) in each sample are responsible for the development of the disease. A recent study estimates the average total number of driver mutations per sample as 4.6, including both SNVs (Single Nucleotide Variants) and CNVs (Copy Number Variants) [31]. Current strategies focus on first identifying a subset of genes that contain driver mutations (‘driver genes’) and then determining which mutations in those genes are drivers. Driver genes are identified on the basis of containing a statistically higher number of cancer somatic mutations than sample background, together with other factors [5,32–40]. Although a number of sets of driver genes have been proposed, there is limited agreement between them [41]. It is very likely that some other genes contain some driver mutations, and conversely it is clear that driver genes will contain some level of non-driver (‘passenger’) mutations. In this work we used the driver gene sets derived from two cancer mutation datasets [6,42].

A number of methods for identifying individual driver missense mutations [43–48] have been developed. These primarily utilize combinations of driver gene lists, predicted impact of mutations, clustering of mutations, and the number of samples in which a mutation has been observed. A limited number of missense driver mutations have been reliably annotated, for example, a set of 889 (Catalog of Validated Oncogenic Mutations, CVOM, https://www.cancergenomeinterpreter.org/mutations [49]). For analysis of mutation properties, these sets may contain significant biases, for example an emphasis on repeat occurrence, favoring oncogene mutations over tumor suppressors. We address the problem of uncertain driver mutation assignments by considering all mutations found in the sets of driver genes, and investigating properties of interest as a function of driver confidence related features.

### 5. Questions addressed

We use the monogenic disease and cancer driver mutation data together with the computational methods to address the following questions:

How effective are sequence-based methods for identifying mutations relevant to the two types of disease? As noted above, these methods depend on mutations impacting Darwinian fitness and, especially for cancer, the validity of that assumption is not clear. Technical issues may also limit accuracy.

How important are intrinsically disordered regions of proteins compared with ordered regions in the two types of disease? The role of disordered regions in protein function has been much discussed [50,51], but the relationship to disease-causing mutations is not yet clear.

What is the relative role of mutations on the protein surface versus those in the core of protein structures? Surface mutations are more likely to be involved in inter-molecular interactions and other mechanisms, while core mutations will be enriched for effects on protein structure stability. How extensive is the role of destabilization of protein structure in the two types of disease? As noted above, this mechanism plays a major role in monogenic disease, but its role in cancer has been less clear.

What are the properties of mutations in cancer passenger genes? Are these benign, as the ‘passenger’ designation implies?

## Results

### 1. Performance of variant interpretation methods on monogenic disease and cancer missense mutations

We begin the analysis by investigating the fraction of monogenic disease mutations and assumed cancer drivers that are predicted to be deleterious by a number of sequence-based methods. As noted earlier, these methods indirectly utilize the impact of a mutation on fitness. Figure 1A shows the results using three different sequence methods (SNPs3D profile [17], Polyphen2 [20] and CADD [19]), comparing the fraction of mutations predicted to be deleterious on a monogenic disease dataset, HGMD [14], and mutations in two sets of cancer driver genes [52–59]. These methods have previously been shown to be effective for identifying monogenic disease mutations [25,60,61], and consistent with that, the fraction predicted deleterious here is high, between 0.85 and 0.89. For all three methods, the fraction deleterious for mutations in cancer driver genes is substantially lower (0.64 - 0.74) (Figure 1A). An additional four monogenic disease missense analysis methods and three developed specifically for cancer missense analysis show the same pattern (Table 1). Previous studies have also shown similar results for cancer data [62,63].

**Table 1.** Performance of sequence-based variant interpretation methods on all datasets

| Variant interpretation method |  | Fraction of predicted deleterious mutations |  |  | Fraction of predicted benign interspecies variants |  |  | Fraction of deleterious cancer missense mutations |  |
| --- | --- | --- | --- | --- | --- | --- | --- | --- | --- |
|  |  | HGMD | TCGA Sander | Cosmic Gene Census | HGMD | TCGA Sander | Cosmic Gene Census | TCGA Passenger | Cosmic Passenger |
| General | SNPs3D Profile | 0.85 | 0.64 | 0.64 | 0.90 | 0.92 | 0.92 | 0.57 | 0.58 |
|  | <sup>a</sup>SNPs3D Profile | 0.84 | 0.64 | 0.64 | 0.90 | 0.92 | 0.91 | 0.57 | 0.58 |
|  | <sup>b</sup>SNPs3D Profile | 0.82 | 0.72 | 0.72 | 0.87 | 0.89 | 0.89 | 0.62 | 0.63 |
|  | PPH2 | 0.89 | 0.74 | 0.70 | 0.94 | 0.97 | 0.97 | 0.64 | 0.64 |
|  | CADD | 0.88 | 0.66 | 0.64 | 0.90 | 0.90 | 0.90 | 0.52 | 0.53 |
|  | SIFT | 0.76 | 0.67 | 0.65 | 0.90 | 0.99 | 0.99 | 0.57 | 0.58 |
|  | LRT | 0.83 | 0.73 | 0.72 | 0.89 | 1.00 | 1.00 | 0.58 | 0.59 |
|  | VEST3 | 0.92 | 0.81 | 0.79 | 0.90 | 0.90 | 0.90 | 0.65 | 0.66 |
|  | PROVEAN | 0.81 | 0.55 | 0.53 | 0.92 | 0.96 | 0.97 | 0.45 | 0.46 |
|  | SNPs3D Stability | 0.72 | 0.49 | 0.47 | 0.87 | 0.86 | 0.88 | 0.41 | 0.42 |
|  | <sup>b</sup>SNPs3D Stability | 0.71 | 0.53 | 0.50 | 0.86 | 0.83 | 0.84 | 0.44 | 0.45 |
| Cancer specific | FATHMM | 0.81 | 0.38 | 0.40 | 0.39 | 0.68 | 0.70 | 0.18 | 0.19 |
|  | Mutation Assessor | 0.74 | 0.48 | 0.45 | 0.95 | 0.97 | 0.97 | 0.40 | 0.42 |
|  | CHASM | 0.43 | 0.49 | 0.61 | 0.90 | 0.90 | 0.90 | 0.24 | 0.22 |

<sup>a</sup>Method retrained separated for disordered and ordered regions
<sup>b</sup>Method retrained separated for protein core and surface regions

**Figure 1:**
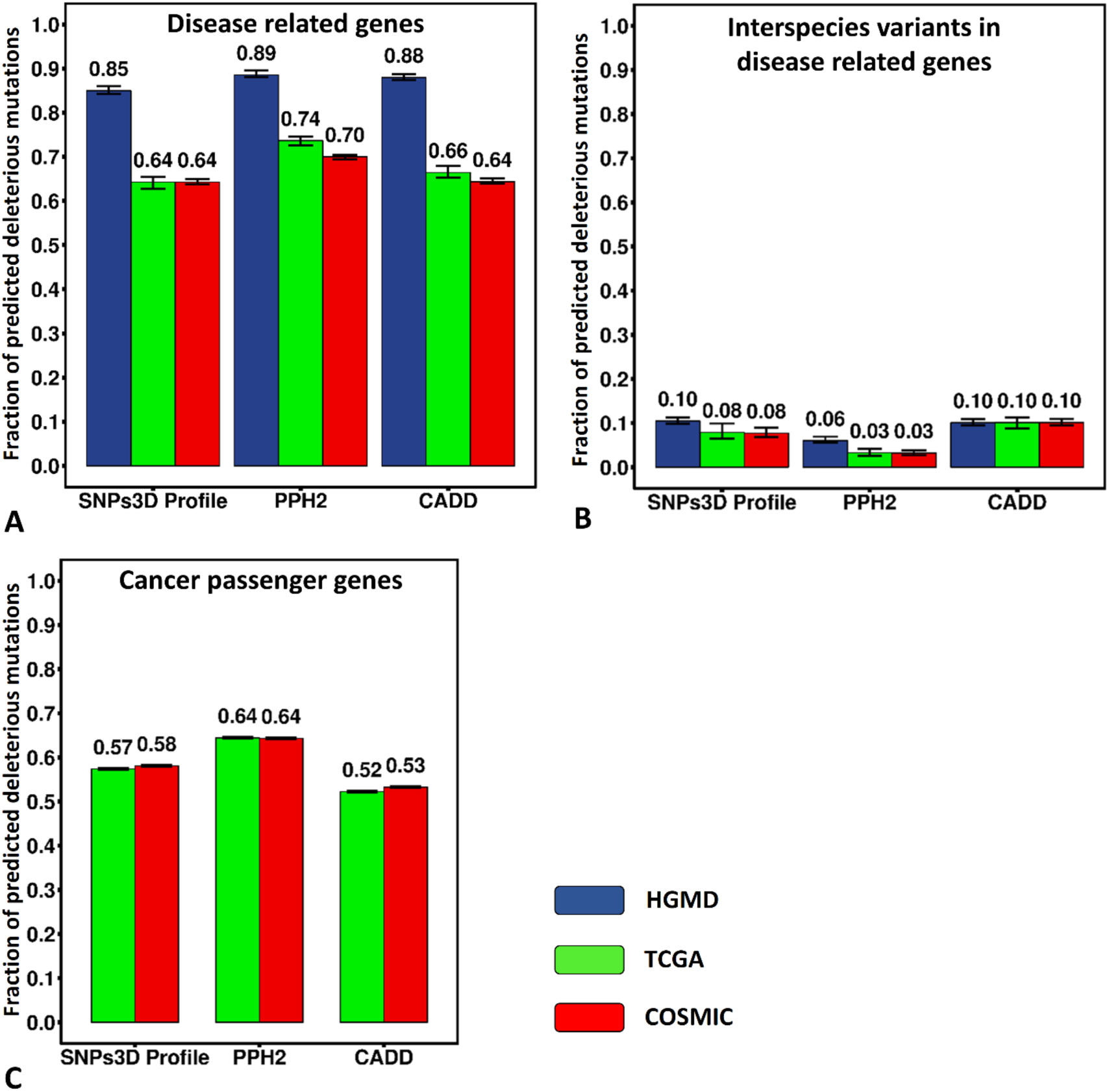
Performance of three sequence-based variant interpretation methods on mutations in the two types of diseases. (**A**) Fraction of predicted deleterious mutations for monogenic disease mutations (HGMD, blue), cancer somatic mutations in the Sander driver gene list (TCGA Sander, green), and cancer somatic mutations in the COSMIC Cancer Gene Census driver gene list (COSMIC, red). A consistently high fraction of monogenic disease mutations appear deleterious, while the fractions of mutations in cancer driver genes are consistently lower. (**B**) Fraction of predicted deleterious interspecies variants in these gene sets. A very low fraction is predicted deleterious in all three data sets, supporting a low false positive rate for the methods. (**C**) Fraction of predicted deleterious mutations in passenger genes in the two cancer data sets. A surprisingly high fraction of mutations are predicted deleterious compared with the interspecies controls, suggesting limited purifying selection and the presence of additional driver mutations. Error bars show 95% confidence intervals derived from 100 rounds of bootstrapping. PPH2 is Polyphen-2.

Figure 1B shows that in contrast to the results for the disease mutations, interspecies variants in these three sets of genes have a uniformly low predicted deleterious fraction (0.03 - 0.10), with no significant differences between monogenic disease and cancer. Although a small number of human monogenic disease mutations have been found to be fixed in other species [64], these numbers do provide an approximate measure of the false positive rate for the methods.

Surprisingly, for mutations in passenger genes (Figure 1C) the predicted deleterious fraction is much higher (0.52 – 0.64) than for the interspecies variants in the driver genes. As discussed later, there are two possible factors contributing here: weak purifying selection in cancer samples (suggested by others [65]), and additional drivers in genes currently designated as passengers. Supplementary Table 1 provides the number of missense mutations and other statistics for all datasets.

### 2. The effect of passenger mutations in cancer driver genes

A likely reason for the lower fraction of mutations predicted deleterious for cancer is that not all mutations in driver genes are drivers, for example, the mutations within the C-terminus in the APC protein [5]. We used two methods to define subsets of mutations enriched in drivers. The first assumes that the more cancer samples a mutation is observed in, the more likely it is to be a driver, an approach that others have also used [6]. Figure 2 shows the dependence of the predicted deleterious fraction (PDF) on the number of occurrences of a mutation. Strikingly, the PDF in driver genes increases sharply with mutation recurrence, from approximately 0.6 for mutations only observed once to 0.9 for those observed more than 10 times. The latter value is higher than the PDF for monogenic disease. For mutations in passenger genes, on the other hand, the PDF does not increase with recurrence and is lower than the lowest value for driver genes. Thus, by this criterion, the low PDF observed in the driver gene mutation set is primarily a consequence of the presence of passenger mutations in driver genes, and a pure driver set would have a PDF values as high as or higher than that for monogenic disease.

**Figure 2:**
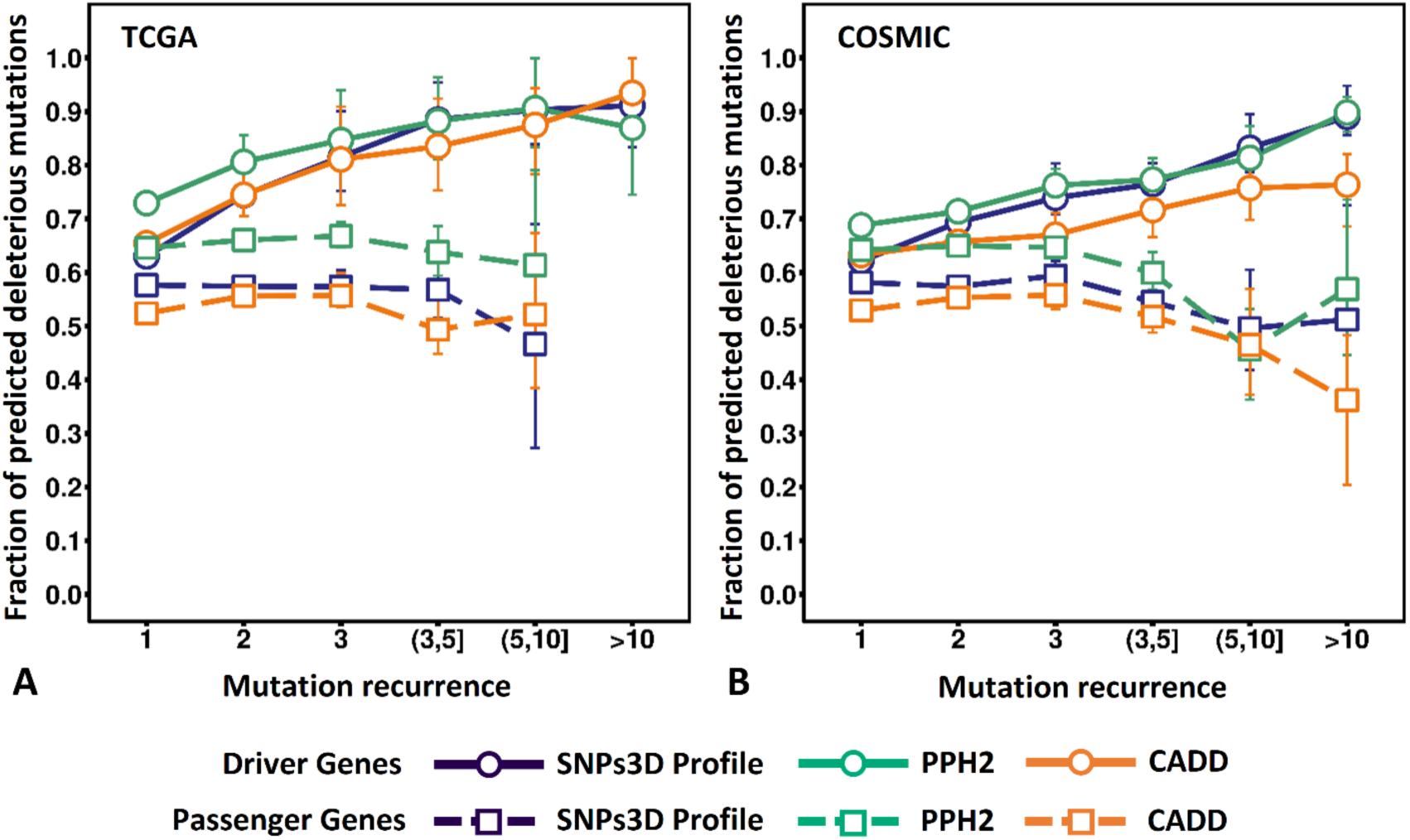
Fraction of predicted deleterious mutations in driver genes (circles and solid lines) as a function of mutation recurrence, for two cancer datasets. The fraction rises from around 0.6 for single occurrence mutations to about 0.9 for those occurring more than 10 times. In contrast to that, for passenger gene mutations (squares and dashed lines), the value is approximately constant for all recurrence values, and lower than for the lowest recurrence driver gene value. These data are consistent with an increase in the fraction of true driver mutations with recurrence. For the most enriched driver set, the fraction predicted deleterious is higher than that for monogenic disease (**Figure 1**). The results are consistent across SNPs3D Profile (blue), PPH2 (Polyphen-2, green) and CADD (orange). Error bars show 95% confidence intervals derived from 100 rounds of bootstrapping.

The second method of enriching for driver mutations examines the PDF as a function of the average total mutational load in different cancer types. As noted above, mutational load differs by more than two orders of magnitude, depending on cancer type [2,5], so that the background of passengers in low mutation load cancers will be very much smaller than for high load ones. Thus, if passengers in driver genes are a cause of the low overall PDF, the PDF will be higher in cancer types with a lower mutational load. Figure 3 shows that this is the case: for driver genes, the trend is for increased PDF as mutation load decreases, whereas passenger genes show no trend. These results are consistent across three different variant interpreting methods (Supplementary Figure 1). Thus, by this criterion too, the low PDF observed in the driver gene mutation set is primarily a consequence of the presence of passenger mutations in driver genes.

**Figure 3:**
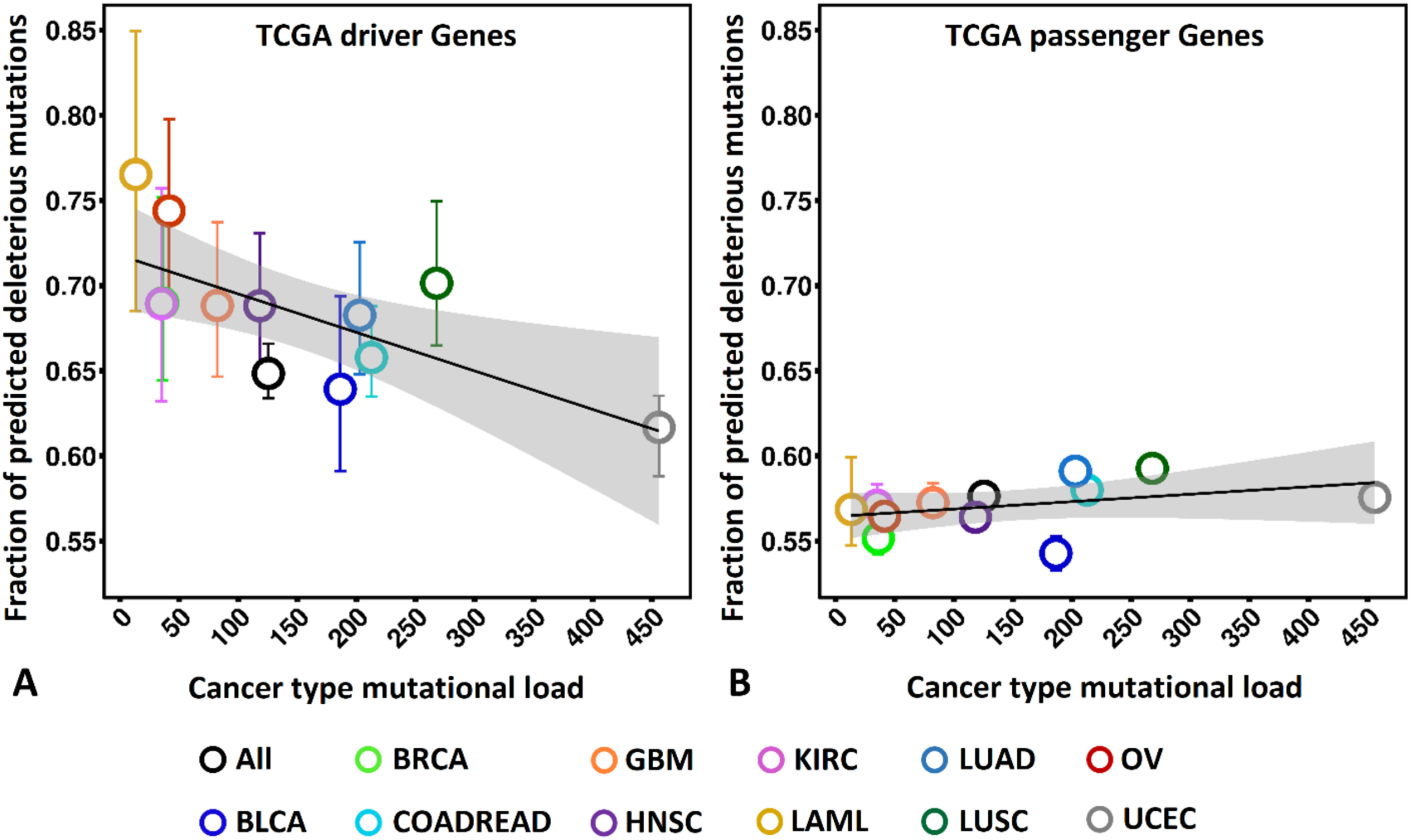
Fraction of predicted deleterious mutations as a function of average mutational load for different cancer types. **(A)** The fraction of predicted deleterious mutations in TCGA Sander driver genes is negatively correlated with the mutation load across cancer types, whereas (B) the fraction of predicted deleterious mutations in the TCGA passenger genes does not show correlation with the mutation load. The correlation is consistent across SNPs3D Profile (shown here), PPH2 (shown in **Supplementary Figure 1**) and CADD (**Supplementary Figure 1**). The results are consistent with the overall low predicted deleterious fraction arising from the burden of passenger mutations in driver genes. Error bars show 95% confidence intervals inferred from 100 rounds of bootstrapping. Small error bars may be obscured by symbols. BLCA: Bladder urothelial carcinoma; BRCA: Breast invasive carcinoma; COADREAD: Colon and rectum adenocarcinoma; GBM: Glioblastoma multiforme; HNSC: Head and neck squamous cell carcinoma; KIRC: Kidney renal clear-cell carcinoma; LAML: Acute myeloid leukemia; LUAD: Lung adenocarcinoma; LUSC: Lung squamous cell carcinoma; OV: Ovarian serous cystadenocarcinoma; UCEC: Uterine corpus endometrioid carcinoma.

### 3. Other factors that may affect the fraction of driver gene mutations predicted deleterious

We explored two other possible explanations for the different deleterious rates for monogenic disease and cancer mutations. One difference between the two types of disease is that whereas most monogenic disease missense mutations overwhelmingly result in loss of protein function [25], cancer driver mutations are either loss of function (in tumor suppressors) or gain of function (in oncogenes). The cancer data were divided into these two subtypes and the analysis repeated (Table 2). Results vary a little by method, but overall there is no substantial difference between the two types of driver genes, so this is not a significant factor in the monogenic disease/cancer difference. A second possible explanation is training bias - methods trained on one type of disease may not perform as well on the other. Two lines of evidence show this is also not a significant factor. First, the analysis done with the three cancer-specific methods shows a similar monogenic disease/cancer difference (Figure 1, Table 1). Second, versions of the SNPs3D Profile method retrained on the cancer datasets also produce a lower predicted deleterious fraction in cancer genes than in monogenic disease (Supplementary Table 2 shows statistics on this and all other retraining results).

**Table 2.** Performance of variant interpretation methods on cancer oncogene and tumor suppressor gene subsets

| Variant interpreting method |  | Fraction of predicted deleterious mutations in oncogenes |  | Fraction of predicted deleterious mutations in tumor suppressor genes |  |
| --- | --- | --- | --- | --- | --- |
|  |  | TCGA Sander | Cosmic Gene Census | TCGA Sander | Cosmic Gene Census |
| General | SNPs3D Profile | 0.66 | 0.68 | 0.75 | 0.70 |
|  | <sup>a</sup>SNPs3D Profile | 0.65 | 0.67 | 0.75 | 0.70 |
|  | <sup>b</sup>SNPs3D Profile | 0.67 | 0.73 | 0.80 | 0.74 |
|  | PPH2 | 0.72 | 0.72 | 0.77 | 0.73 |
|  | CADD | 0.79 | 0.68 | 0.71 | 0.66 |
|  | SIFT | 0.69 | 0.67 | 0.75 | 0.68 |
|  | LRT | 0.87 | 0.82 | 0.73 | 0.75 |
|  | VEST3 | 0.88 | 0.83 | 0.87 | 0.83 |
|  | PROVEAN | 0.60 | 0.59 | 0.60 | 0.57 |
|  | SNPs3D Stability | 0.47 | 0.44 | 0.59 | 0.53 |
|  | <sup>b</sup>SNPs3D Stability | 0.51 | 0.47 | 0.61 | 0.55 |
| Cancer specific | FATHMM | 0.37 | 0.39 | 0.52 | 0.57 |
|  | MutationAssessor | 0.52 | 0.45 | 0.53 | 0.49 |
|  | CHASM | 0.75 | 0.80 | 0.71 | 0.80 |

<sup>a</sup>Method retrained separated for disordered and ordered regions
<sup>b</sup>Method retrained separated for protein core and surface regions

### 4. Intrinsically disordered regions in monogenic disease and cancer

We next examine the first of three protein structure related factors that affect the properties of monogenic disease and cancer mutations: the role of intrinsically disordered structure. 27% of residues in monogenic disease proteins are predicted disordered by the method used here [66] (Figure 4A). Cancer passenger proteins, representing the majority of genes, have a similar value (Figure 4A). But for cancer driver proteins, as others have also noted [67], the predicted content of disordered residues is substantially higher, at 41 to 45% (Figure 4A). (The disorder data are derived using a machine learning prediction method [66] rather than direct observation of structure, 5% false positive rate threshold).

**Figure 4:**
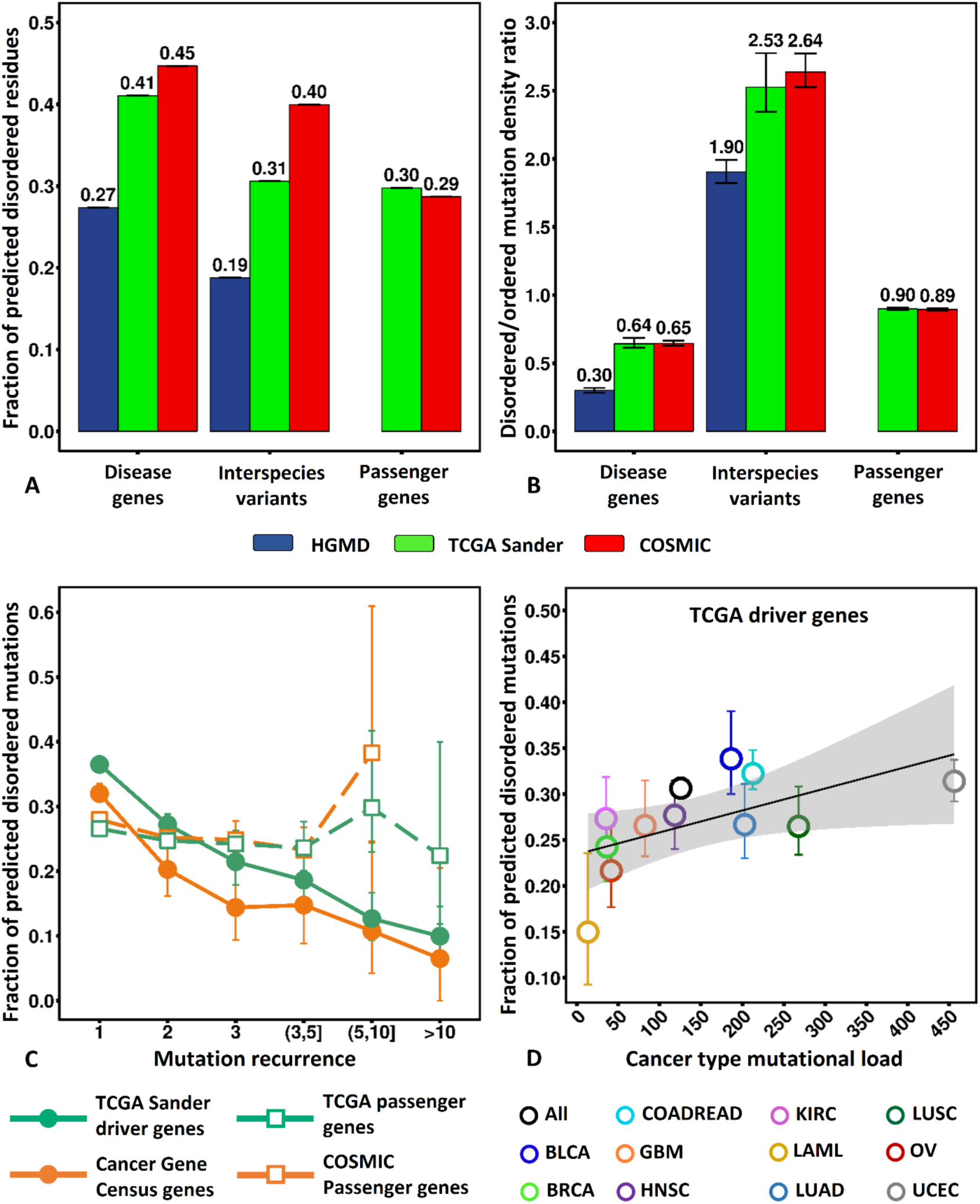
(**A**) Predicted fraction of intrinsically disordered residues. Only about ¼ of monogenic disease protein residues are predicted disordered (blue), compared with nearly twice as many in cancer driver genes (green and red). Passenger gene values are close to those for monogenic disease. (**B**) Ratio of mutation density in disordered regions to that in ordered regions. The relative density in cancer driver proteins is more than twice as high as for monogenic disease proteins. High passenger protein relative density and very high values for interspecies variants are consistent with lower functional restraints in disordered regions. (**C**) The fraction of mutations in disordered regions of cancer driver proteins decreases with mutation recurrence rate, consistent with most mutations in these regions being passengers. No dependence on recurrence rate is seen for the equivalent mutations in passenger genes. (**D**) The fraction of mutations in disordered regions of cancer driver proteins increases with cancer type mutational load, also consistent with most of these mutations being passengers. Error bars show 95% confidence intervals derived from 100 rounds of bootstrapping. Small error bars may be obscured by symbols. BLCA: Bladder urothelial carcinoma; BRCA: Breast invasive carcinoma; COADREAD: Colon and rectum adenocarcinoma; GBM: Glioblastoma multiforme; HNSC: Head and neck squamous cell carcinoma; KIRC: Kidney renal clear-cell carcinoma; LAML: Acute myeloid leukemia; LUAD: Lung adenocarcinoma; LUSC: Lung squamous cell carcinoma; OV: Ovarian serous cystadenocarcinoma; UCEC: Uterine corpus endometrioid carcinoma.

### 5. Mutations in intrinsically disordered regions

Figure 4B shows that monogenic disease mutations are only ~1/3 as likely to occur at disordered positions as ordered ones. This, together with the low fraction of disordered residues in the relevant proteins, results in a total of only 10% of monogenic disease mutations lying in disordered regions, indicating a small role for these in this type of disease. In contrast to this, cancer driver gene mutations are only moderately less likely in disordered regions than in ordered ones (0.64~0.65 of the mutation density in ordered regions), and that, together with the higher content of disorder in cancer driver proteins, results in total 31~34% of these mutations occurring in disordered regions. Two factors may contribute to the higher disorder cancer mutation density - an excess of passenger mutations in disordered regions, and a possible greater functional role for disordered regions in cancer drivers than in monogenic disease.

Figure 4B also shows that the density of mutations in the ordered and disordered regions of passenger proteins is approximately equal and slightly higher than that of driver proteins. The highest relative density is for interspecies variants, with a density more than 2.5 times higher in disordered regions than ordered ones. Both this and the higher passenger relative density are consistent with substantially less functional restrictions on the acceptance of mutations in disordered regions and so a tendency for passengers to accumulate there. In support of this, Figure 4C shows that the fraction of driver gene mutations observed more than 10 times (and therefore most likely to drivers) in disordered regions is only 1/3 the fraction for mutations observed only once (so least likely to be drivers). Similarly, Figure 4D shows that the fraction of mutations in disordered regions decreases with decreasing total mutational load, consistent with a higher fraction of passengers in these regions. Conservatively, these results indicate the maximum fraction of driver mutations in disordered regions is about 25%. Part of this excess over monogenic disease protein disordered regions is due to the larger fraction of disorder residues in the cancer proteins - about 1.5 fold higher. Thus, it does not appear that there is a much larger role of mutations in disordered regions for cancer.

### 6. Fraction of deleterious mutations in ordered and disordered regions

We next examine predicted deleterious rates in disordered versus ordered regions, using the sequence methods introduced earlier. Since all sequence methods are trained on a full set of disease mutations, and, particularly for monogenic disease, there are more mutations in ordered than disordered regions, training bias was a concern here. To investigate the extent of bias, we trained versions of the SNPs3D profile method on only disordered HGMD mutations together with disordered interspecies variants as controls and also trained on the corresponding ordered data. In fact, retraining made almost no difference to the results: the predicted deleterious fraction (PDF) in ordered regions is unchanged at 0.85. The original PDF in disordered regions is 0.86 and for the retrained method is 0.84. Correction for a second factor does have a significant impact on the results. Inspection of the set of HGMD mutations in disordered regions revealed that a substantial fraction (438 out of the total of 1110) are in collagen. At first glance, it seems odd that collagen should be classed as a disordered protein, but this is a correct characterization - the bulk of collagen molecules are formed from a homo-triple helix. A hypothetical monomer would be structurally disordered, and the repeat triplet of the sequence is one of the possible signatures of disordered regions. Nevertheless, from the point of view of this analysis, the collagen mutations are atypical of disordered regions in other proteins, so we again retrained the SNPs3D profile method, omitting these mutations. The PDF in disordered regions, omitting the collagen mutations, is now substantially lower (0.69 versus 0.84). Figure 5A shows that result on monogenic disease mutations together with those from Polyphen2 [20] and CADD [19], omitting the collagen mutations. The results from all three methods are similar and show consistently lower PDFs for disordered versus disordered regions (0.85 - 0.89 in ordered regions, 0.66-0.69 in disordered regions). That is, for monogenic disease, the fraction of mutations predicted deleterious is about 20% lower for disordered than ordered regions. The reason for this is unclear.

**Figure 5:**
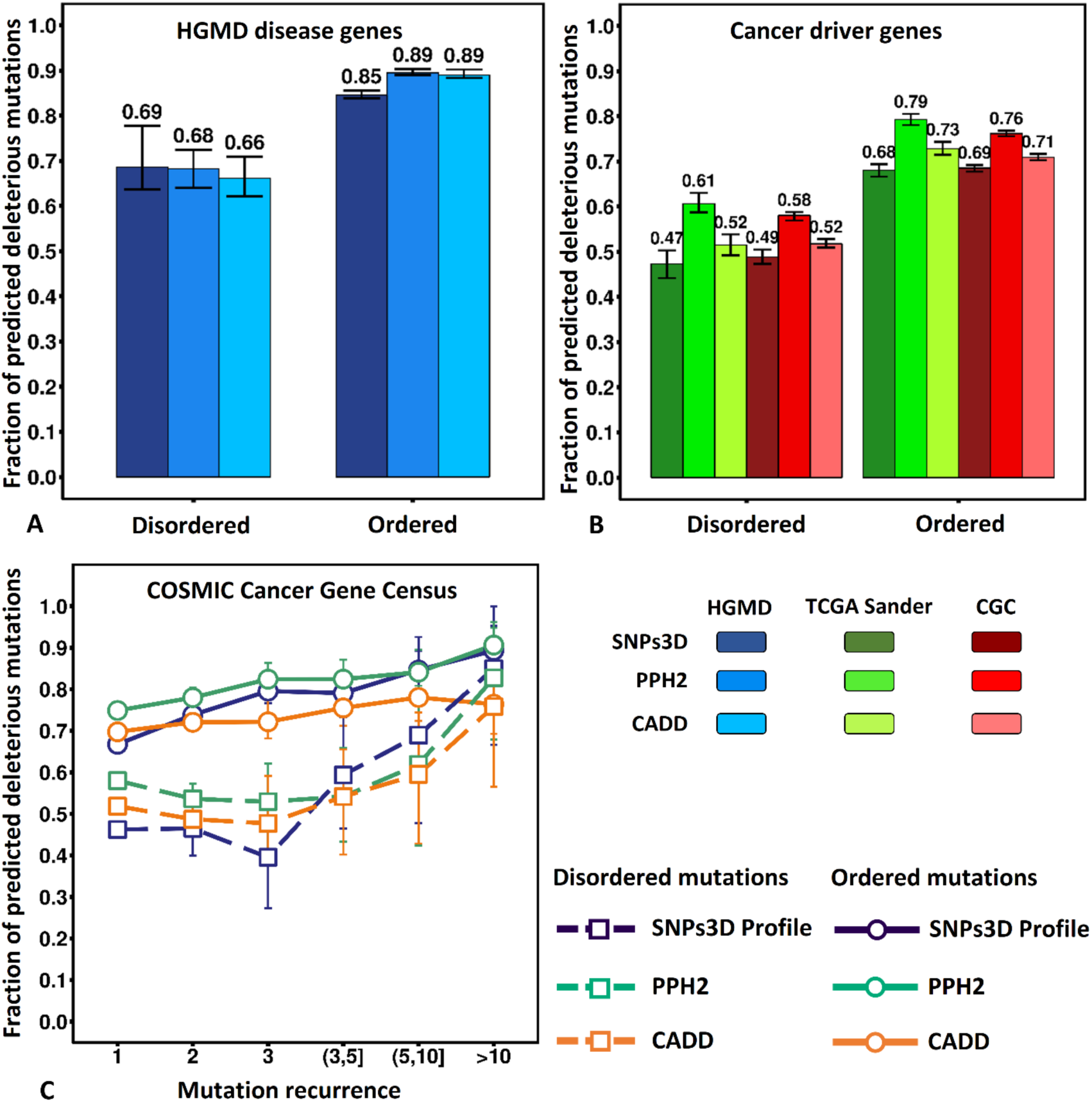
(**A**) The fraction of predicted deleterious mutations is approximately 20% lower in disordered regions of monogenic disease proteins than in ordered regions. (**B**) For cancer driver proteins, the fraction of predicted deleterious mutations in disordered regions is approximately 30% lower than for ordered regions. (**C**) Fraction of predicted deleterious mutations in the ordered (circles and solid lines) and disordered (square and dashed lines) regions of COSMIC Cancer Gene Census driver proteins as a function of mutation recurrence. For mutations with low recurrence, the fraction of predicted deleterious mutations is consistently lower in disordered regions than in ordered regions. Both fractions rise as a function of mutation recurrence and converge when mutations are observed for more than 10 times. Error bars show 95% confidence intervals derived from 100 rounds of bootstrapping.

Figure 5B shows that the fraction of cancer driver gene mutations predicted deleterious in disordered regions is also consistently about 20% lower than in ordered regions.

### 7. Other properties of mutations in disordered versus ordered regions

We found no tendency for mutations in tumor suppressors and oncogenes to be differently distributed between disordered and ordered regions. Similarly, we found no tendency for monogenic disease mutations in genes classified as dominant versus recessive to be differently distributed in disordered and ordered regions.

### 8. Protein surface and core mutations

A second structural feature that provides insight into the mutation properties is the fraction of mutations on the surface of proteins versus in the core. Figure 6A shows that the fraction of all residues designated ‘surface’ (using STRIDE [68–70], see Methods) is similar for all categories of protein, at approximately 50% (See Figure 7A for the mutations in the core). As shown in Figure 6B, the relative density of monogenic disease mutations on the surface to that in the core is only 0.58, showing a strong tendency for mutations in this class of disease to be buried (See Figure 7B for the mutations in the core). In contrast to this, both cancer driver gene sets show an enrichment of 1.3 for mutations on the surface versus in the core. Passenger gene mutations show a larger surface enrichment of 1.75 on average, and by far the highest surface enrichment is for inter-species variants, 3.6 to 4.2. The latter values reflect the fact that there are more possible neutral mutations on the surface than in the interior, so substitutions are more likely to be fixed on the surface, and there are more opportunities for benign passenger mutations there as well. We therefore expect some of the higher relative density on the surface of driver proteins arises from the accumulation of passengers. However, unlike the data for disordered regions, the fraction of driver gene mutations on the surface as a function of mutation recurrence does not show a significant trend (Figure 6C) and so does not support an excess of surface passengers. Interpretation of this plot is complicated by a strong tendency for oncogene mutations to be on the surface and tumor suppressors to be in the core (see below). Since oncogene mutations have a higher recurrence rate than tumor suppressors, that tendency will dampen any relationship between surface and recurrence. (A plot using the alternative method for assessing the effect of passengers, surface density versus mutation load, does show the expected relationship, but because of limited structural data, 95% confidence limits are large - data not shown). An alternative probe of the extent of passengers in surface regions is to consider the fraction of surface mutations predicted deleterious as a function of recurrence, since this fraction is similar for tumor suppressors and oncogenes (Supplementary Table 2). Figure 6D shows a strong trend of increasing deleterious rate with mutation recurrence, consistent with the results from the passenger proteins and interspecies variant density results. Overall, the data support the conclusion that a substantial part of the excess surface mutations in cancer driver proteins are passenger mutations.

**Figure 6:**
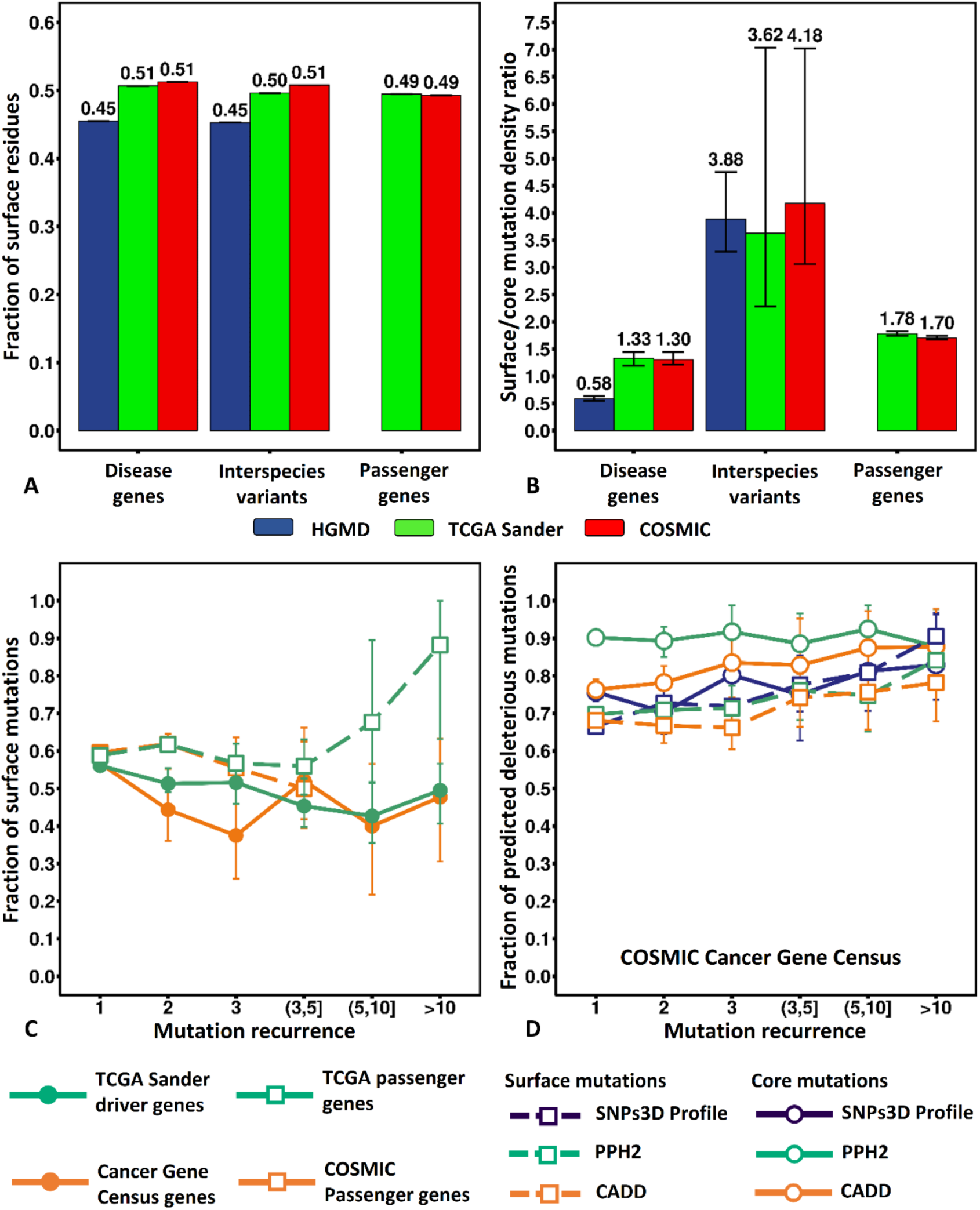
(**A**) For all classes of protein, about half of all residues are designated surface. (**B**) Ratio of mutation density on the surface to that in the core. The density of monogenic disease mutations on the surface is only about ½ that in the core, whereas for cancer driver protein mutations the density is higher on the surface. High ratios for interspecies variants and passenger proteins are consistent with less functional constraints on surface residues. (**C**) The fraction of surface mutations in cancer driver genes does not significantly correlate with mutation recurrence, so does not support an excess of surface mutations being passengers. A confounding factor is the tendency for oncogene mutations to be on the surface. (**D**) Fraction of predicted deleterious mutations in the core (circles and solid lines) and on the surface (square and dashed lines) for COSMIC Cancer Gene Census driver proteins as a function of mutation recurrence. The fraction of predicted deleterious mutations on the surface rises from around 0.7 for single occurrence mutations to 0.8~0.9 for those occurring more than 10 times. Error bars show 95% confidence intervals derived from 100 rounds of bootstrapping.

**Figure 7.**
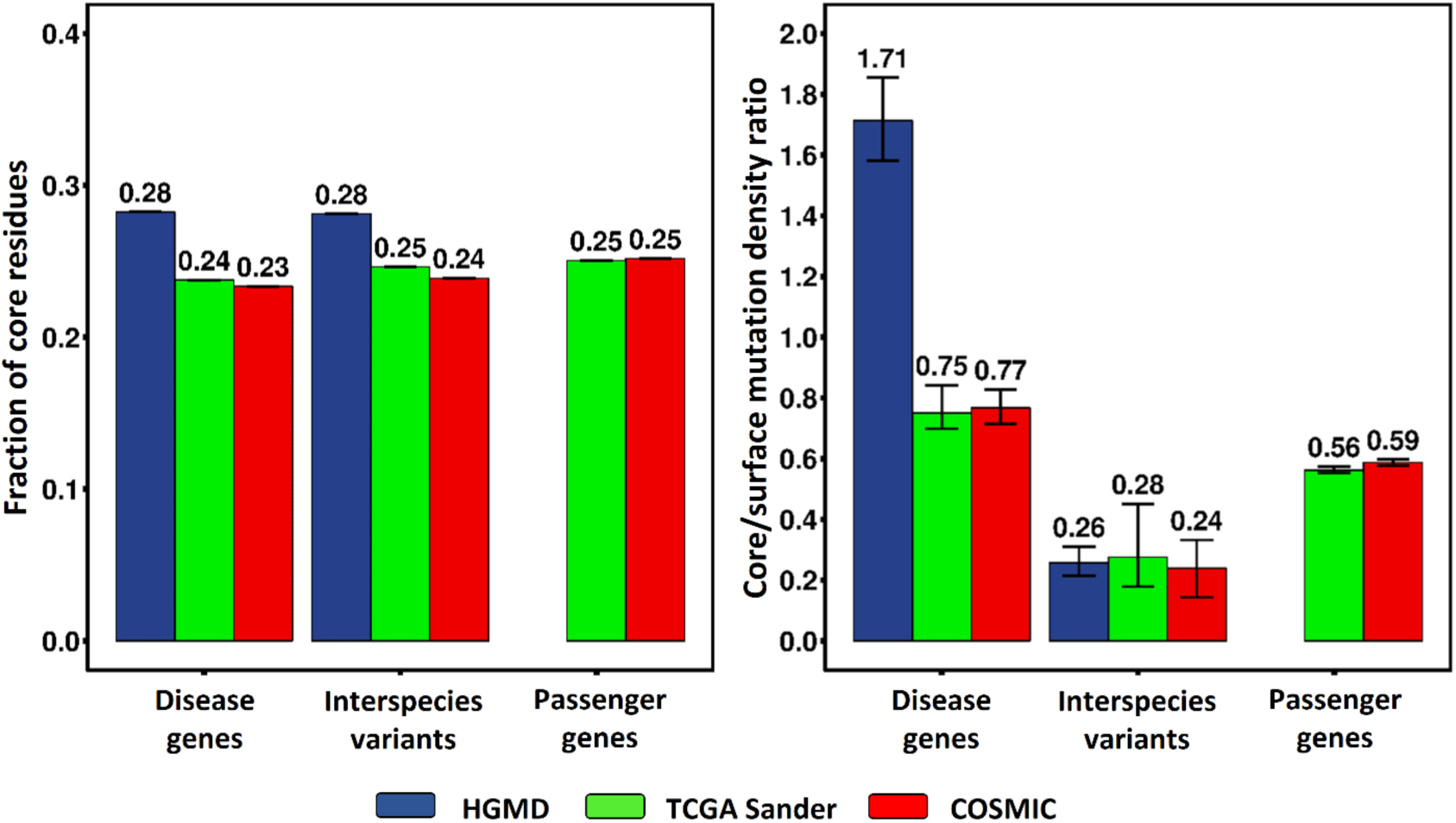
(A) Fractions of core residues and (B) ratio of mutation density in the core to that on the surface in monogenic disease genes and cancer driver genes, in the corresponding interspecies variants datasets, and in cancer passenger genes. The density ratio in HGMD disease genes is significantly higher than in the cancer driver genes. Compared to **Figure 5**, interspecies variants and somatic mutations in the passenger genes are more enriched on the protein surface, which supports that surface missense mutations are less deleterious and more tolerated.

We also examined the surface to core distribution for mutations in oncogenes and tumor suppressors (Supplementary Figure 2). Unlike the corresponding data for disorder/order, there is a marked difference in surface enrichment for the two classes of genes: for oncogenes the density of surface mutations is 1.9 times that of core mutations, while for tumor suppressors, the density is lower on the surface than in the core (average 0.85 that of the core). For monogenic disease, there is also a smaller but still significant difference between the surface to core densities for genes classified as dominant and those classified as recessive, (density ratio of 0.68 for dominant versus 0.45 for recessive).

### 9. Role of structure destabilization

We next examine the role of structure destabilization in cancer mutations compared to those in monogenic disease. As noted earlier, destabilization plays a major role in monogenic disease mechanisms [25] and our earlier analysis suggests a significant role in cancer too [26]. For this purpose, we trained separate stability SVMs on surface and core monogenic disease mutations, with interspecies variants in those regions as controls. Figure 8A shows that overall 0.71 of monogenic disease mutations are predicted to be destabilizing (similar to the value Yue & Moult reported earlier [17]), whereas only about 0.50 to 0.53 cancer driver gene mutations are predicted destabilizing, a large difference. The interspecies variant results provide an estimated false positive rate of 0.15. To address possible bias arising from training the stability method on monogenic disease data, we retrained using cancer data. This model was unsatisfactory in that it delivered much higher false positive rates (0.24 to 0.41 of interspecies variants predicted destabilizing, Supplementary Table 2), but the relationship between the fraction of monogenic disease mutations predicted destabilizing and the fraction for cancer driver genes is similar to that obtained with the monogenic disease model (0.81-0.84 for monogenic disease and 0.64-0.68 for cancer driver gene mutations, Supplementary Table 2), supporting the conclusion that destabilization rates are substantially higher for monogenic disease than for cancer driver gene mutations.

**Figure 8:**
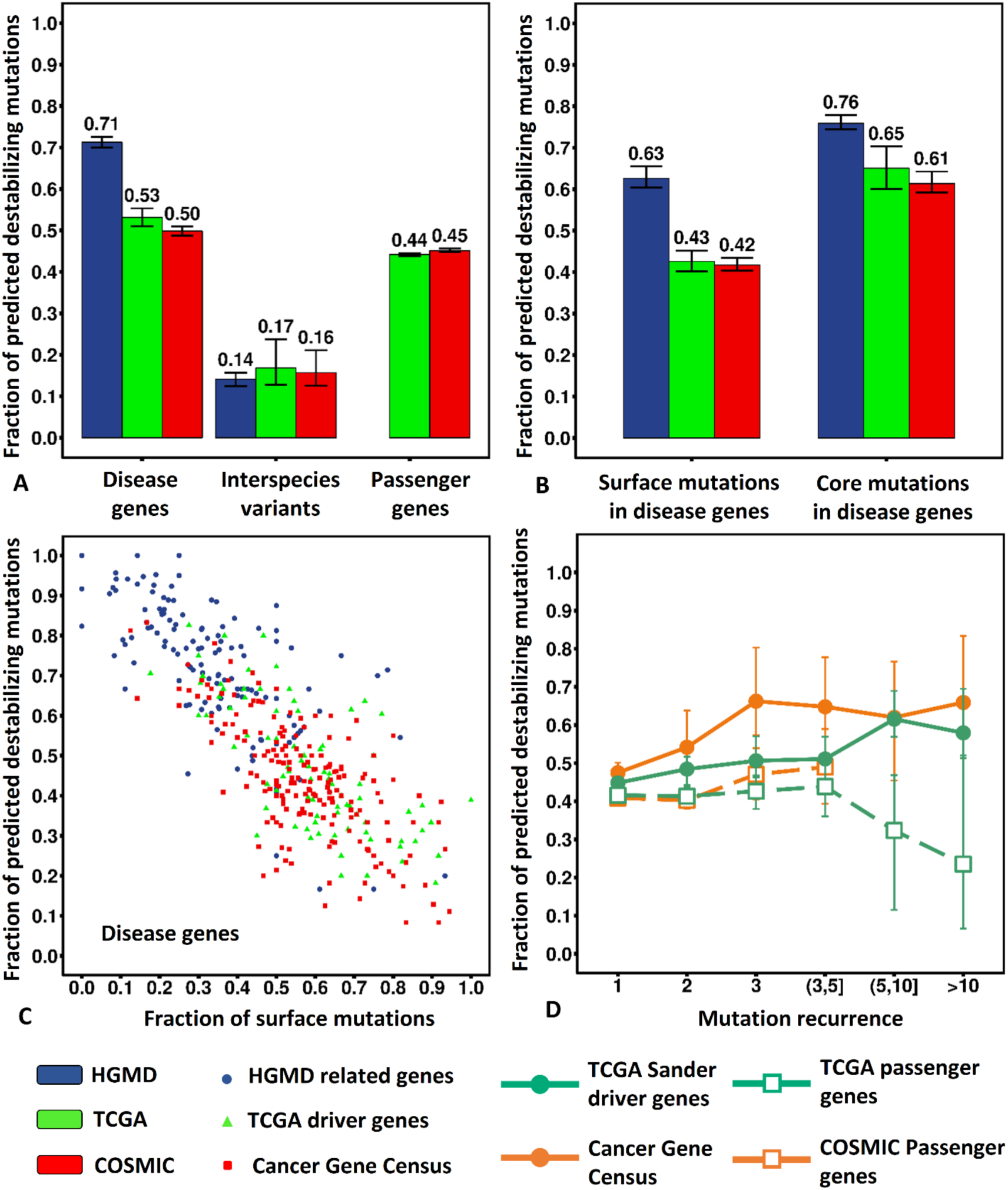
Fraction of predicted destabilizing mutations. (**A**) More than 70% of monogenic disease mutations are predicted destabilizing compared with only about half of mutations in cancer driver genes. The value for passenger gene mutations is not much lower than for driver genes. (**B**) The predicted destabilizing fraction is substantially lower on the protein surface than in the core for both types of disease. (**C**) Dependence on the fraction of mutations predicted destabilizing on the fraction of surface mutations. Each point is for one gene. The fewer mutations on the surface, the higher the fraction predicted destabilizing, and monogenic disease genes (blue) tend to have a lower surface fraction than cancer genes. (**D**) Relationship between the fraction predicted deleterious and recurrence for cancer mutations. The driver gene destabilizing fraction increases with recurrence, consistent with a mixture of driver and passenger mutations in these genes. There is no trend for passenger gene mutations. Results are for SVMs trained on monogenic disease surface and core mutations separately. Error bars show 95% confidence intervals derived from 100 rounds of bootstrapping.

Based on the other analyses, we expect that passenger mutations in driver genes are one contributor to the lower fraction predicted destabilizing there. Examination of the fraction of destabilizing mutations as a function of mutation recurrence (Figure 8D) supports a role for this factor, although less strongly than in the corresponding sequence analysis. A second contribution to different levels of destabilization in cancer and monogenic disease may come from the higher proportion of surface mutations in cancer: surface mutations are intrinsically less likely to be destabilizing. To help isolate this effect, we examined the role of destabilization in surface and core mutations separately. As expected, values for surface and core are markedly different: For monogenic disease, 0.76 of mutations in the core are predicted destabilizing, compared with 0.63 for surface. For cancer driver gene mutations, average values are 0.63 for core and 0.42 for surface (Figure 8B).

We expected that the distinction between surface and core destabilization properties might be particularly sensitive to whether a mutation is an oncogene or a tumor suppressor, so also examined the surface/core properties of these two classes separately. Indeed, tumor suppressor destabilizing fractions are higher in the core (~0.70) than those for oncogenes (~0.56), while surface values are similar for the two classes of mutation (Supplementary Table 2, Supplementary Figure 2). Thus for tumor suppressors and monogenic disease mutations, a high (>70%) fraction of core mutations destabilize protein structure.

## Discussion

In this paper, we have used computational methods together with sequence and structure information to investigate and compare the properties of missense mutations causative of monogenic disease and those driving cancer. The principal findings are as follows:

### 1. Most monogenic disease and cancer driver mutations are under selection pressure, and so can be identified with sequence-based methods

After allowing for the effects of passenger mutations in cancer driver genes, we find a high fraction (> 80%) of mutations causing monogenic disease and of cancer driver mutations are predicted to be deleterious. These results are consistent across three different methods trained on monogenic disease (Figures 3, Supplementary Figure 1). There are two primary implications. First, sequence methods trained on monogenic disease data are effective at identifying cancer drivers. Second, the large majority of mutations positions in both types of disease are under strong selection pressure (otherwise the sequence methods used would not be effective). While this was likely for most monogenic diseases, which are often severe and early onset, it is less obvious for cancer mutations, selected within a clone primarily to promote cell growth. It is not yet clear what fraction of the apparent false negatives - disease mutations not predicted deleterious - are technical false negatives or mutations that do not affect fitness.

### 2. Mutations in disordered regions play a limited role in both monogenic disease and cancer

Figure 4A shows that there is a much higher involvement of disordered regions in cancer than in monogenic disease, while Figure 4B shows that the relative density of driver protein mutations in these regions is twice as high as for mutations in monogenic disease genes (~0.64 versus 0.30). Together, this leads to only 10% of monogenic disease mutations in these regions, compared with 31-34% of cancer driver protein mutations. Two factors contribute to this difference. First, the even higher relative mutation density in passenger protein disordered regions (~0.9) and for interspecies variants (~2.6) suggests that part of the cancer excess density is a consequence of benign passenger mutations being more likely to lie in disordered regions. Analysis of the relative densities as a function of the mutation recurrence and cancer mutational load confirm this is the case. The fraction of mutations predicted deleterious in disordered regions of cancer driver proteins is also low compared with that for monogenic disease, also consistent with a high fraction of passengers in these regions (Figure 5). Second, cancer driver genes are unusual in containing almost 1.5 times as much disorder as monogenic disease genes or passenger genes. That higher disordered fraction likely reflects a different functional spectrum for these proteins. In particular, it has been noted that these proteins are more hub-like [71–73] (involved in interactions with many partners), perhaps implying that more disordered regions are required to provide specificity for a range of protein binding partners [74,75]. For example, the intrinsically disordered terminal trans-activation domain of P53 binds to three different protein partners in three different conformations [76]. As noted above, the apparently deleterious mutations in disordered regions may often be involved in protein-protein interactions. However, the extent to which disordered regions of proteins are involved in function has not been clear [50]. Indeed, the two aspects of these results confirm that disordered regions are much less functionally significant - the very low fraction of monogenic disease mutations there, and the high concentration of interspecies variants and passenger mutations.

### 3. Cancer oncogene mutations tend to be on the protein surface, whereas monogenic disease mutations and tumor suppressor mutations tend to be in the core

The surface density of mutations in cancer driver genes is higher than in the core, and for mutations in oncogenes, it is nearly twice as high. While some of this difference reflects excess passenger mutations on the surface, it also likely reflects the greater role for disruption of intermolecular interactions in cancer [77] and also that gain-of-function oncogene mutations tend to affect surface processes such as kinase conformational states related to phosphorylation [78]. Conversely, tumor suppressor mutation density is higher in the core than on the surface, and for monogenic disease mutations, the core density is twice that of the surface. As discussed below, these values reflect the large role of structure destabilization for these classes of mutation. In monogenic disease, there is a higher relative density of surface mutations in autosomal dominant genes than recessive ones (0.68 versus 0.45). A number of mechanisms are involved in autosomal dominant disease, including haplo-insufficiency, oligomer structure (of which collagen mutations are an example [79]), and gain of function. The latter mechanism likely contributes most to the surface/core signal, in a manner analogous to that of gain of function mutations in oncogenes. Examples of monogenic disease surface gain-of-function mutations are for the calcium sensing receptor (CASR), causing hypocalcemia, and for Luteinizing Hormone/Choriogonadotropin Receptor (LHCGR), causing Familial Male-Limited Precocious Puberty (FMPP).

### 4. A large fraction of monogenic disease and cancer tumor suppressor mutations in the protein core destabilize protein structure

Approximately ¾ of both monogenic disease mutations and cancer tumor suppressor mutations in the protein core are predicted to destabilize protein structure. A prediction of destabilization using this method is equivalent to a major decrease in protein abundance *in vivo* [25]. Most monogenic disease missense mutations result in a major loss of molecular function (for example, [81] and [61]). To the extent that core tumor suppressor mutations can be considered to represent all driver mutations, the result implies that in this class of disease too there is usually major loss of protein function, rather than a subtle effect at that level.

There are some oncogene mutations in the core region, and about 50% of these are predicted to destabilize protein structure, at first glance a surprising result, since these should be gain-of-molecular function. As noted earlier, less than 1/3 of oncogene mutations are in the core, so that a 50% destabilization rate corresponds to just 1/6 of oncogene mutations. The estimated false positive rate is 0.15, close to that value, and there may be some cases where oncogene gain of function is the result of destabilization of a regulatory domain. Also, the definition of oncogenes and tumor suppressors is not always unambiguous. In compiling the oncogene and tumor suppressor lists we noted that 10 genes had been classified as oncogenes by one group and tumor suppressors by the other (these were excluded). There are also examples where a gene may behave as an oncogene in some circumstances and a tumor suppressor in others [82].

### 5. Consistent results on the curated cancer driver mutation data

As noted earlier, conclusions about cancer driver mutations depend on methods that can distinguish these from passenger mutations. To check the generality of our results, we applied similar analyses to the 613 TCGA and CGC mutations included in the 889 missense mutations in the Catalog of Validated Oncogenic Mutations (CVOM) (https://www.cancergenomeinterpreter.org/mutations, [49]). This is a small manually selected dataset and so should contain few passenger missense mutations. 84% of mutations in this subset are predicted deleterious by SNPs3D Profile, a similar number to that obtained for recurring mutations in our analysis, consistent with the recurrence bias in this dataset. The CVOM set is biased towards higher recurrence mutations (25% mutations occurring more than 10 times as opposed to 1% in the full TCGA driver set and 2% in the CGC driver set). As observed for our high recurrence mutations (Figure 4C) a very high fraction (93%) of these CVOM mutations are in ordered regions, consistent with the conclusion that passenger mutations in cancer driver genes are likely enriched in disordered regions. The CVOM set is also biased towards oncogene mutations (84%), hence is dominated by surface mutations surface (79%), again consistent with our findings. The fraction of CVOM core mutations predicted to destabilize protein structure in tumor suppressors is 0.9. Counts here are small, but the result is consistent with the two cancer driver data sets (~0.7), supporting the dominant role of destabilization in these genes. The fraction of destabilizing surface mutations is similar to those in TCGA and CGC driver set (0.37 versus ~0.43),consistent with a smaller role of destabilizing on the cancer driver protein surface.

### 6. A high fraction of mutations in passenger genes are deleteriousness

A surprisingly high fraction (more than 0.5, Supplementary Figure 1 and Table 1) of mutations in passenger genes appear deleterious with the sequence methods used here. The estimated false positive rate is much lower (10% or less). A more conservative measure using consensus among seven distinctive sequence based methods (requiring at least 5 out 7 methods to predict deleterious) led to similar results (0.45~0.46). Stability analysis supports this observation, with almost as high a fraction of passenger gene mutations predicted destabilizing as in driver genes. There are at least two possible explanations. One is that there is insufficient selection pressure to eliminate these mutations in a typical cancer. Simulations of cancer progression suggest that moderately deleterious mutations will escape elimination by various population genetics mechanisms, and so accumulate, sometimes impending cancer progression [65]. Still, this is a high fraction. Using the same method on missense population SNPs (greater than 1% population frequency in dbSNP137) yields a value of about 34% (similar results are obtained with other methods (Supplementary Table 3)). The other is that there is a significant concentration of unrecognized driver genes. As noted earlier, there is considerable variation in driver set definitions, so that it is expected this to some degree. But depending on the cancer type and particular case [2], there may be up to 100s or even a thousand deleterious mutations spread across non-driver genes. A striking difference of the passenger gene predicted deleterious mutations compared to those in driver genes is that the fraction predicted deleterious is largely independent of the number of observations (Figure 2). The reason for this is not obvious. The exact nature and impact of these mutations will repay further study.

In common with all analyses so far, we have assumed a binary model of cancer drivers - a mutation is either a driver or not. But it may be that there is a continuous scale of driver impact with a few strong drivers and a long tail of mutations making secondary contributions, loosely analogous to the contributions of variants to complex trait disease [83].

## Materials and Methods

### 1. Monogenic disease and cancer missense mutation data

The monogenic disease set comprises 10,865 disease-related variants collected from an earlier version of HGMD [14], together with 13,499 interspecies variants in these genes, compiled by comparing mammalian homolog protein sequences with at least 90% sequence identity over at least 80% of the full length and excluding any known disease-related variants [17]. The disease genes were classified as dominant or recessive based on inheritance patterns.

Two cancer driver data sets were compiled as follows. One set was extracted from the level 3 TCGA [52–58] data described in [6], which has 449,672 unique somatic missense single-residue substitutions in a total of 3,416 tumor samples from studies on 12 different cancer types (Table 3). The TCGA driver set consists of the 9,325 unique somatic missense mutations found in 193 driver genes identified by [6]. Another 415,090 somatic missense mutations in genes not belonging to the 193 driver gene list were extracted to form the TCGA passenger set. Mutations in other potential driver genes [2,5,29,41,84] were also omitted in the passenger set. 27 oncogenes and 47 tumor suppressor genes were identified in the TCGA 193 driver gene list, based on the literature [5,41,84], providing a TCGA Oncogene set of 1,362 missense mutations and TCGA TSG set of 2,933 missense mutations. A set of 3,116 interspecies variants in the 193 TCGA driver genes were extracted using the same procedure as for the monogenic disease set described above.

The second cancer data set was extracted from the 531,728 unique somatic missense single-residue substitutions in the COSMIC Database [59] version 68. The Cosmic Gene Census (CGC) driver dataset consists of the 30,773 missense mutations in 477 driver genes identified by the Cancer Gene Census [42]. Another 495,530 missense mutations extracted from the COSMIC non-CGC genes form the CGC passenger set. Mutations in other potential driver genes were removed in the same way as for the TCGA passenger set. A CGC Oncogene set of 7,422 missense mutations in 79 genes and a CGC TSG set of 12,016 missense mutations in 81 genes was compiled using the same procedure as for the TCGA sets. A CGC interspecies variant set of 6448 missense mutations was extracted using the same procedures as above.

**Table 3.** TCGA data set

| Tumor type | TCGA ID | Number of unique mutations<sup>a</sup> in driver genes |
| --- | --- | --- |
| Bladder urothelial carcinoma | BLCA | 435 |
| Breast invasive carcinoma<sup>b</sup> | BRCA | 482 |
| Colon and rectum adenocarcinoma<sup>c</sup> | COADREAD | 2170 |
| Glioblastoma multiformae<sup>d</sup> | GBM | 586 |
| Head and neck squamous cell carcinoma | HNSC | 885 |
| Kidney renal clear-cell carcinoma | KIRC | 503 |
| Acute myeloid leukemia<sup>e</sup> | LAML | 140 |
| Lung adenocarcinoma | LUAD | 946 |
| Lung squamous cell carcinoma<sup>f</sup> | LUSC | 939 |
| Ovarian serous cystadenocarcinoma<sup>g</sup> | OV | 460 |
| Uterine corpus endometrioid carcinoma<sup>h</sup> | UCEC | 2223 |

<sup>a</sup>Restricted to somatic missense single-residue substitutions observed in samples of each specific cancer types.
<sup>b</sup>Reference see [52]
<sup>c</sup>Reference see [57]
<sup>d</sup>Reference see [53]
<sup>e</sup>Reference see [56]
<sup>f</sup>Reference see [58]
<sup>g</sup>Reference see [54]
<sup>h</sup>Reference see [55]

### 2. Missense mutation analysis methods

Seven sequence-based missense analysis methods were used to assign missense mutations as deleterious or benign and the fraction of those mutations that are assigned as deleterious (the PDF, predicted deleterious fraction) was calculated for each. Four of these (SNPs3D Profile [17], PolyPhen-2 [20], CADD [19], VEST3 [18,21]) were trained on monogenic disease mutation datasets (except CADD). The other three: SIFT [23], LRT [85], and PROVEAN [86] rely on direct measures of sequence conservation properties and do not require training. In addition, three sequence methods trained specifically for interpreting cancer mutations were tested: FATHMM [48], Mutation Assessor [44] and CHASM [43,87]. SNPs3D Profile results were generated using standalone in-house software. The dbNSFP2.9 database [88] was used to obtain PolyPhen-2, CADD, SIFT, LRT, VEST3, PROVEAN, FATHMM and Mutation Assessor results. CHASM results were obtained from the CRAVAT Web server (http://www.cravat.us/CRAVAT/). Binary assignments of structure destabilizing/non-destabilizing were obtained using SNPs3D Stability [25] with in-house software and the predicted destabilizing fraction (PDF) were calculated from those data.

Binary predictions were collected for PolyPhen-2, SIFT, LRT, PROVEAN, FATHMM and Mutation Assessor. The HumDiv version of PolyPhen-2 was used, and “probably damaging” and “possibly damaging” predictions were considered deleterious. MutationAssessor “H” and “M” predictions were also considered deleterious. Three methods (CADD, VEST3, and CHASM) reported continuous scores rather than binary assignments. Dataset-specific score thresholds were chosen for these, such that the false positive rates on the corresponding interspecies variants sets are similar to that of other methods. For the monogenic disease data, the score thresholds are 22 for CADD, 0.5545 for VEST3, and 0.095 for CHASM. On the TCGA data, the thresholds are 21.35 for CADD, 0.2815 for VEST3, and 0.1395 for CHASM. On the Cosmic data, the thresholds are 21.35 for CADD, 0.2915 for VEST3, and 0.1225 for CHASM.

To assess potential training bias, the SNPs3D Profile and SNPs3D Stability methods were retrained on the two cancer data sets and on specific subsets of monogenic disease data. In retraining, all parameters in the support vector machine (SVM) models were re-optimized with a grid search algorithm. At each search step, the corresponding data set was bootstrapped 30 times, with the model trained on a set of randomly drawn data (number of data points equal to the data set size), and evaluated on the data points not included in training. 95% confidence intervals in the other analyses were also inferred from 30 rounds of bootstrapping.

### 3. Structure modeling

For analysis of structure-related features, the set of experimental protein structures was extended by building homology models for protein domains where a suitable template was available, as described in [25]. The procedure is briefly summarized here. Proteins that have >40% sequence identity to the query protein and a crystal structure of <3Å resolution are used as templates for backbone conformations. The 40% sequence identity cutoff is based on earlier benchmarking [25] that showed prediction accuracy for models based on this or higher sequence identity to a template is not significantly lower than for that based on experimental structures. Where the template amino acids are identical to the corresponding ones in the query structure, side chains atoms from the template are used. Otherwise, the side-chains are modeled using SCWRL [89].

### 4. Analysis of somatic missense mutation recurrence and load

For each unique cancer somatic missense mutation, the recurrence was calculated as the number of times the mutation was observed in all samples. The majority of unique somatic missense mutations, even in the likely driver genes, have a low recurrence (<5). Recurrence values were grouped into six bins, the last of which covers all recurrence larger than 10. The cancer type specific mutation load was defined as the average number of unique missense mutations per sample, observed in the samples of the corresponding cancer type.

### 5. Analysis of structure disorder and surface missense mutations

DISOPRED3.16 [66] with default parameters was used to predict intrinsically disordered protein residues in each data set. STRIDE [68–70] was used to calculate the absolute solvent accessible surface area (SASA) of each amino acid residue in the protein structures or homology models prepared as described in [25] and above. The relative SASA was then calculated by normalizing the STRIDE results with the corresponding amino acid residue maximal solvent accessibility reported in [90]. Based on the relative SASA, residue location was assigned as buried core (<0.05), partially exposed (≥0.05, ≤ 0.25), and surface (>0.25). The relative density (*RD*) of missense mutations in a particular state is calculated as follows:

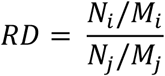

where, given two particular states *i* and *j* (disordered, ordered, buried in the core, and exposed on the surface), *N*_*i*_ and *N*_*j*_ are the total number of missense mutations in the corresponding states, and *M*_*i*_ and *M*_*j*_ are the total number of amino acid residues in the corresponding states.

## Acknowledgements

We thank Lipika Ray Pal and Kunal Kundu for many stimulating discussions and critical feedback. The work was partially support by NIH R01GM120364 to JM.

---

**Supplementary Figure 1:**
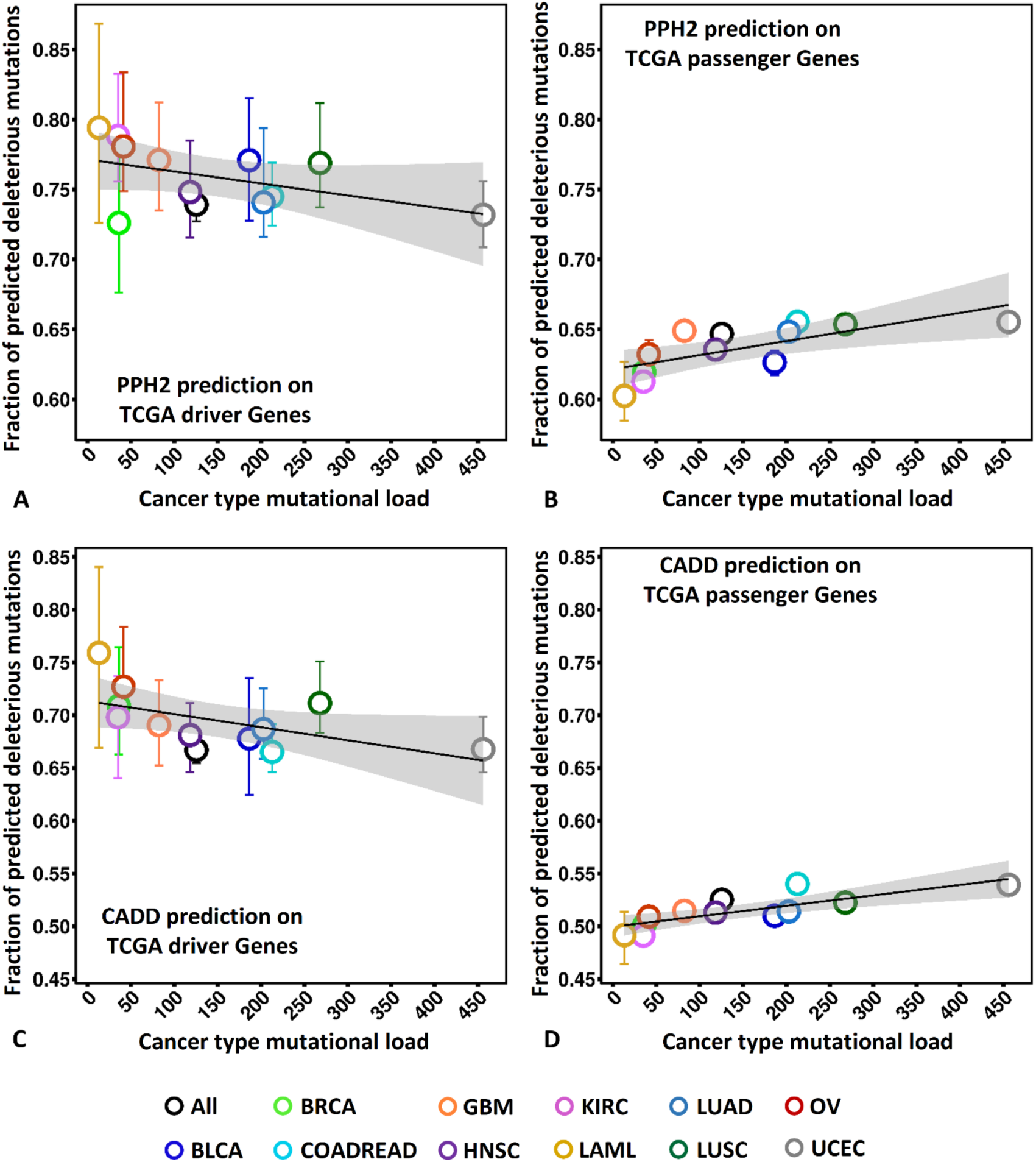
The fraction of predicted deleterious mutations obtained with (A) PPH2 and (C) CADD in driver genes in the TCGA Sander list are negatively correlated with the mutation burden across cancer types, whereas the fraction of predicted deleterious mutations in the TCGA passenger genes (**B, D**) show no or weak positive correlation with the mutation burden. These results are consistent with those from SNPs3D Profile (**Figure 3**). Error bars indicate the 95% confidence intervals inferred from 100 round bootstrapping. Small error bars may not be obscured by the symbols. BLCA, Bladder urothelial carcinoma. BLCA: Bladder urothelial carcinoma; BRCA: Breast invasive carcinoma; COADREAD: Colon and rectum adenocarcinoma; GBM: Glioblastoma multiforme; HNSC: Head and neck squamous cell carcinoma; KIRC: Kidney renal clear-cell carcinoma; LAML: Acute myeloid leukemia; LUAD: Lung adenocarcinoma; LUSC: Lung squamous cell carcinoma; OV: Ovarian serous cystadenocarcinoma; UCEC: Uterine corpus endometrioid carcinoma.

**Supplementary Figure 2.**
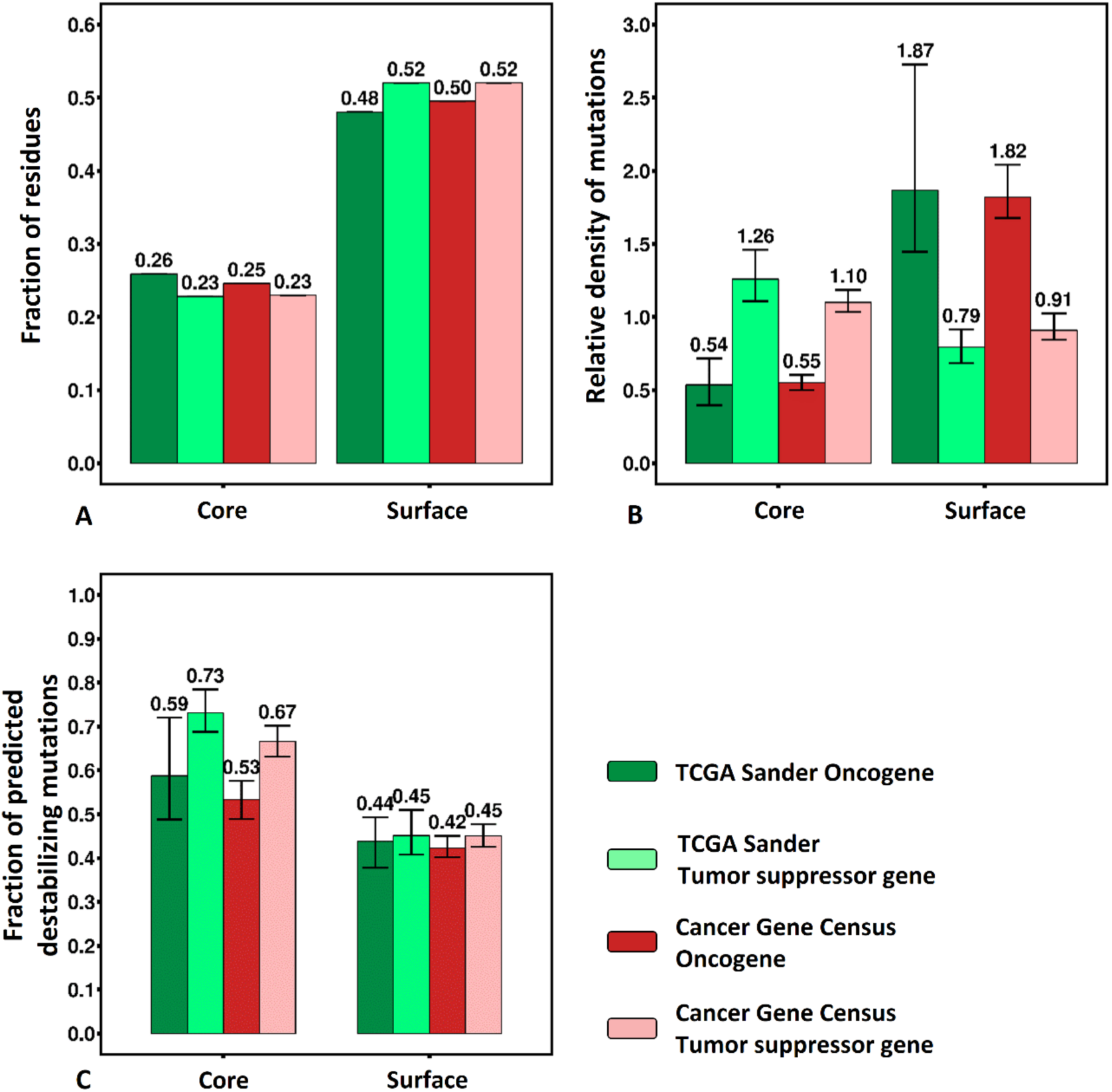
(**A**) The relative residue fraction and (**B**) the relative mutation density, and (**C**) the fraction of mutations predicted destabilizing for core and surface mutations in the TCGA Sander oncogene set, the TCGA Sander tumor suppressor gene set, the COSMIC Cancer Gene Census oncogene set, and the COSMIC Cancer Gene Census tumor suppressor gene set. Missense mutations are enriched in the core in tumor suppressor genes, and on the surface in oncogenes. In the core, the fraction predicted destabilizing is significantly higher in tumor suppressor genes than in oncogenes.

**Supplementary Table 1.** Total number of mutations and proteins (in brackets) in each dataset, and coverage of these by each variant interpretation methods

| Prediction Method | HGMD | HGMD Interspecies | TCGA Sander | TCGA Sander Interspecies | TCGA Passenger | TCGA Sander Oncogene | TCGA Sander Tumor Suppressor | Cosmic Gene Census | Cosmic Gene Census Interspecies | Cosmic Gene Passenger | Cosmic Gene Census Oncogene | Cosmic Gene Census Tumor Suppressor |
| --- | --- | --- | --- | --- | --- | --- | --- | --- | --- | --- | --- | --- |
| **SNPs3D Profile** | 10004<br>(745) | 10121<br>(284) | 6706<br>(203) | 1834<br>(74) | 307829<br>(17185) | 1131<br>(28) | 1970<br>(50) | 22763<br>(546) | 3492<br>(180) | 380489<br>(15860) | 6215<br>(87) | 7859<br>(103) |
| **PPH2** | 10002<br>(727) | 10415<br>(266) | 8412<br>(212) | 2659<br>(77) | 401899<br>(17544) | 1354<br>(29) | 2886<br>(58) | 29540<br>(519) | 5558<br>(187) | 494897<br>(16098) | 7123<br>(86) | 11544<br>(98) |
| **CADD** | 9869<br>(724) | 10351<br>(267) | 8454<br>(214) | 2678<br>(77) | 404224<br>(17621) | 1354<br>(29) | 2887<br>(58) | 29559<br>(520) | 5586<br>(187) | 495491<br>(16139) | 7123<br>(86) | 11545<br>(98) |
| **SIFT** | 4235<br>(502) | 6855<br>(237) | 8379<br>(214) | 2678<br>(77) | 400115<br>(17433) | 1354<br>(29) | 2838<br>(58) | 29276<br>(516) | 5556<br>(185) | 493072<br>(16024) | 7107<br>(84) | 11287<br>(97) |
| **LRT** | 9647<br>(699) | 10028<br>(262) | 7682<br>(174) | 2527<br>(76) | 359929<br>(16210) | 1285<br>(25) | 2783<br>(48) | 28114<br>(507) | 5469<br>(186) | 455807<br>(15292) | 6807<br>(84) | 10772<br>(93) |
| **VEST3** | 10004<br>(723) | 10438<br>(265) | 8405<br>(210) | 2668<br>(77) | 402427<br>(17596) | 1354<br>(29) | 2885<br>(58) | 29554<br>(520) | 5558<br>(187) | 495491<br>(16139) | 7123<br>(86) | 11544<br>(98) |
| **PROVEAN** | 10004<br>(723) | 10438<br>(265) | 8426<br>(213) | 2668<br>(77) | 402391<br>(17590) | 1354<br>(29) | 2887<br>(58) | 29559<br>(520) | 5558<br>(187) | 495322<br>(16138) | 7123<br>(86) | 11545<br>(98) |
| **SNPs3D Stability** | 6096<br>(460) | 1771<br>(194) | 3067<br>(133) | 297<br>(46) | 79764<br>(6810) | 765<br>(28) | 1264<br>(41) | 11431<br>(359) | 447<br>(89) | 100579<br>(6448) | 4058<br>(73) | 4819<br>(80) |
| **FATHMM** | 665<br>(233) | 549<br>(106) | 8361<br>(211) | 2643<br>(75) | 383121<br>(16183) | 1350<br>(29) | 2876<br>(57) | 29148<br>(507) | 5333<br>(181) | 473737<br>(14960) | 7059<br>(86) | 11543<br>(97) |
| **Mutation Assessor** | 657<br>(232) | 547<br>(105) | 8375<br>(200) | 2658<br>(77) | 398147<br>(16957) | 1349<br>(28) | 2860<br>(54) | 29311<br>(513) | 5523<br>(186) | 490181<br>(15705) | 7080<br>(83) | 11449<br>(98) |
| **CHASM** | 10865<br>(784) | 13484<br>(288) | 9325<br>(234) | 3116<br>(82) | 412778<br>(17698) | 1362<br>(30) | 2933<br>(61) | 30767<br>(554) | 6448<br>(197) | 494101<br>(16063) | 7421<br>(90) | 12016<br>(104) |
| **DISOPRED3** | 10865<br>(784) | 13499<br>(288) | 9233<br>(233) | 3116<br>(82) | 410792<br>(17715) | 1362<br>(30) | 2933<br>(61) | 30773<br>(554) | 6448<br>(197) | 493231<br>(16061) | 7422<br>(90) | 12016<br>(104) |
| **<sup>a</sup>Ordered** | 9755<br>(738) | 9371<br>(282) | 6379<br>(215) | 1474<br>(71) | 296136<br>(17414) | 1040<br>(29) | 1935<br>(52) | 20210<br>(537) | 2339<br>(175) | 362346<br>(15924) | 5927<br>(88) | 7563<br>(102) |
| **<sup>a</sup>Disordered** | 1110<br>(261) | 4128<br>(243) | 2854<br>(196) | 1642<br>(77) | 114656<br>(13142) | 322<br>(21) | 998<br>(54) | 10563<br>(490) | 4109<br>(182) | 130885<br>(12459) | 1495<br>(79) | 4453<br>(101) |
| **STRIDE** | 6095<br>(460) | 1722<br>(193) | 3055<br>(132) | 297<br>(46) | 79704<br>(6800) | 765<br>(28) | 1263<br>(41) | 11428<br>(359) | 447<br>(89) | 100504<br>(6441) | 4057<br>(73) | 4818<br>(80) |
| **<sup>b</sup>Core** | 2156<br>(302) | 199<br>(96) | 590<br>(93) | 30<br>(17) | 13568<br>(4354) | 122<br>(22) | 309<br>(33) | 1272<br>(252) | 39<br>(24) | 17799<br>(4391) | 643<br>(59) | 1112<br>(65) |
| **<sup>b</sup>Surface** | 2025<br>(352) | 1244<br>(187) | 1676<br>(127) | 219<br>(45) | 47645<br>(6630) | 435<br>(26) | 586<br>(41) | 6209<br>(350) | 346<br>(86) | 59308<br>(6311) | 2283<br>(73) | 2330<br>(78) |

<sup>a</sup>Ordered or disordered mutation subsets predicted by DISOPRED3
<sup>b</sup>Core or surface mutation subsets from STRIDE

**Supplementary Table 2.**
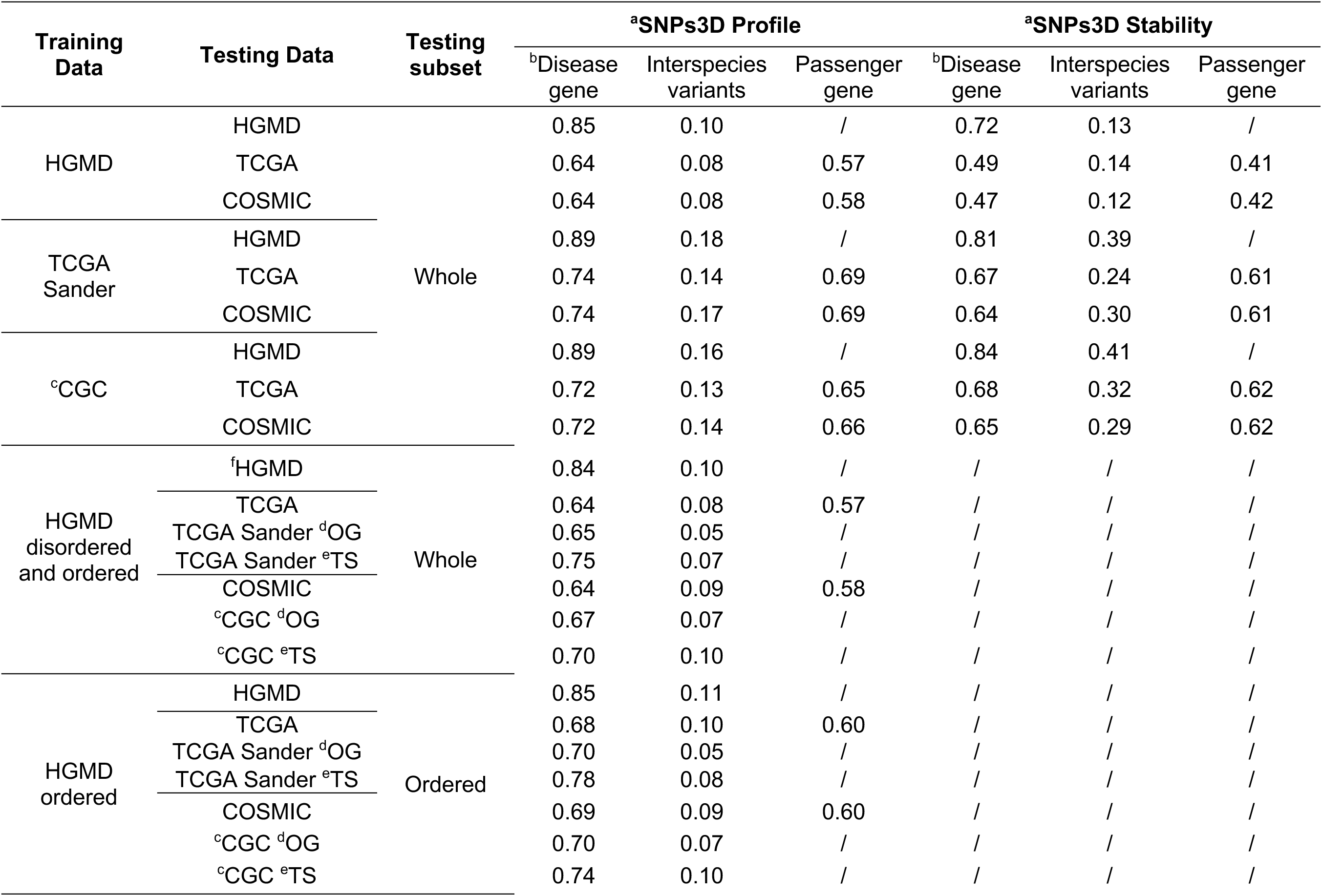
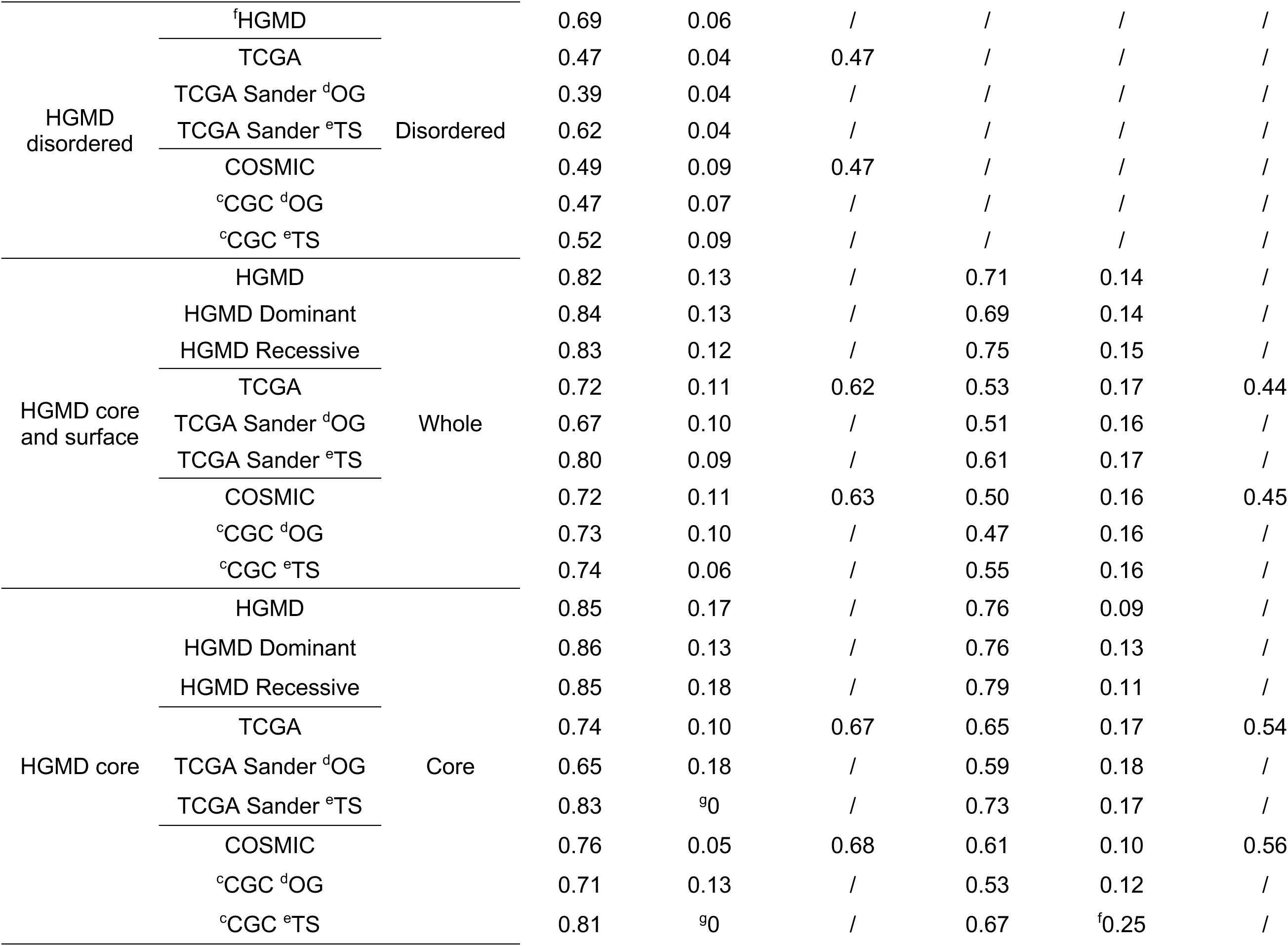
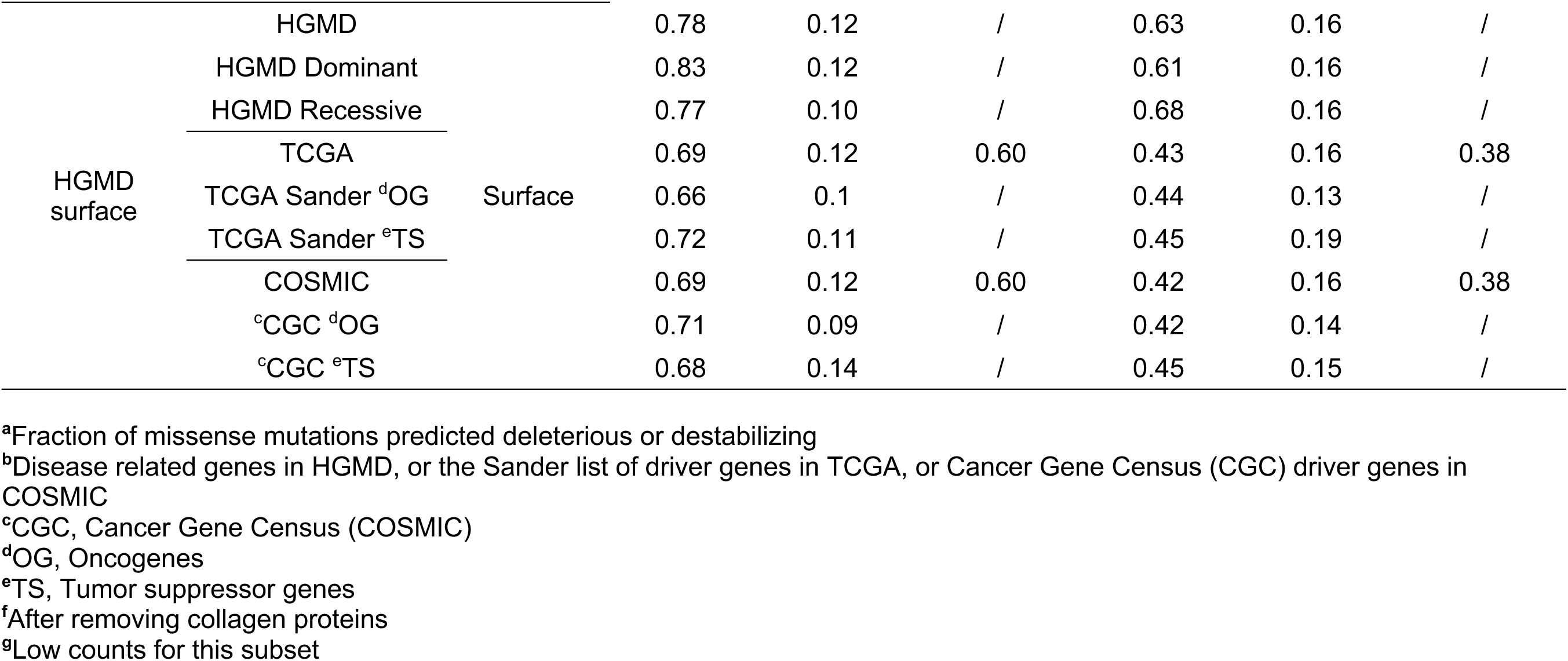
Performance of SNPs3D methods trained on specific datasets

**Supplementary Table 4.**
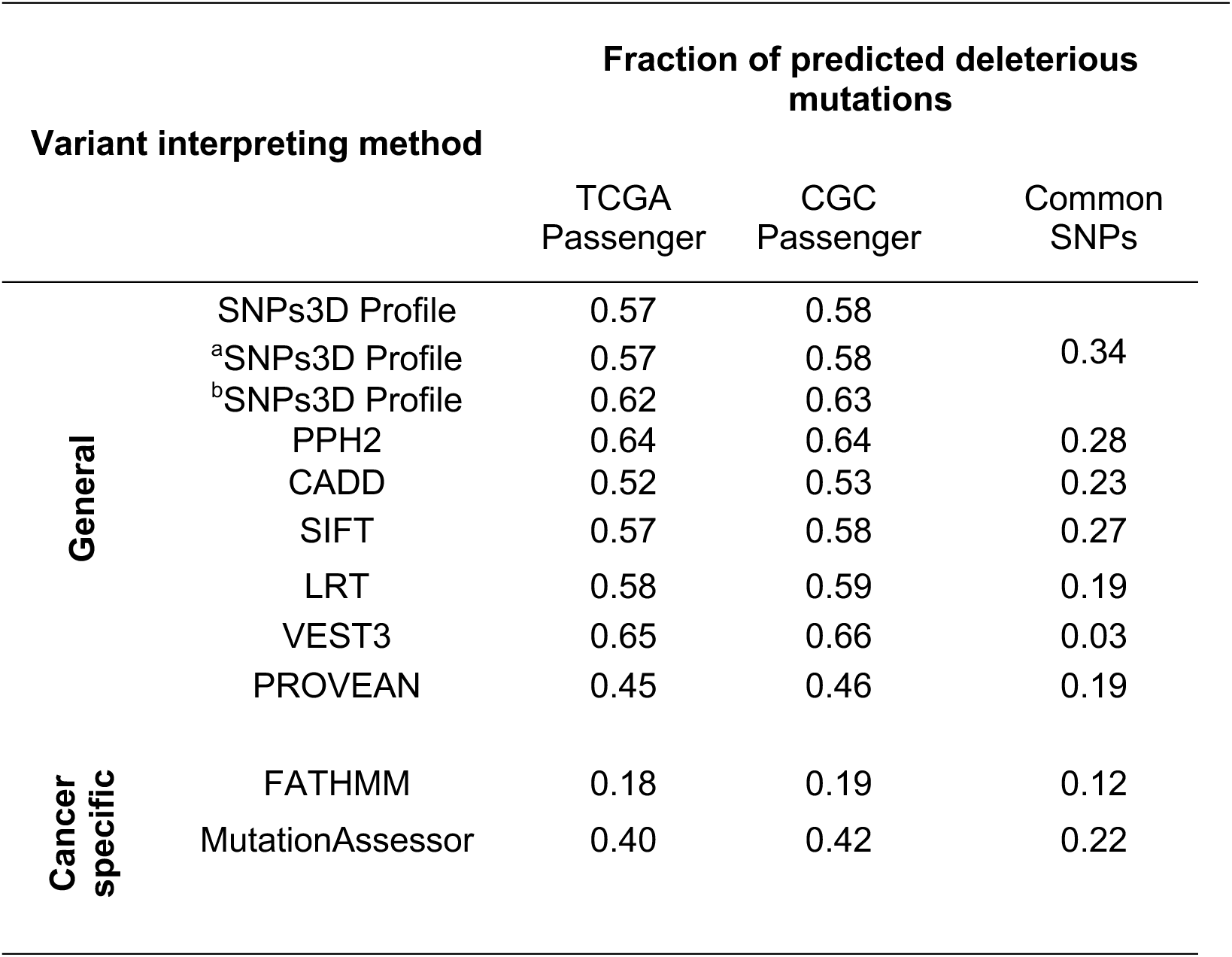
Predicted fractions of deleterious missense mutations in cancer passenger genes and in missense common SNPs (>1%)

## Notes

Grant Sponsor This work was supported in part by NIH R01GM120364 to JM.

## References

[1] J. Shendure, J.M. Akey, The origins, determinants, and consequences of human mutations., Science. 349 (2015) 1478–1483.

[2] I. Martincorena, P.J. Campbell, Somatic mutation in cancer and normal cells., Science. 349 (2015) 1483–1489.

[3] P.D. Stenson, M. Mort, E. V. Ball, K. Evans, M. Hayden, S. Heywood, M. Hussain, A.D. Phillips, D.N. Cooper, The Human Gene Mutation Database: towards a comprehensive repository of inherited mutation data for medical research, genetic diagnosis and next-generation sequencing studies, Hum. Genet. 136 (2017) 665–677.

[4] M.J. Landrum, J.M. Lee, M. Benson, G. Brown, C. Chao, S. Chitipiralla, B. Gu, J. Hart, D. Hoffman, J. Hoover, W. Jang, K. Katz, M. Ovetsky, G. Riley, A. Sethi, R. Tully, R. Villamarin-Salomon, W. Rubinstein, D.R. Maglott, ClinVar: public archive of interpretations of clinically relevant variants., Nucleic Acids Res. 44 (2016) D862–868.

[5] B. Vogelstein, N. Papadopoulos, V.E. Velculescu, S. Zhou, L.A. Diaz, K.W. Kinzler, Cancer genome landscapes., Science. 339 (2013) 1546–1558.

[6] G. Ciriello, M.L. Miller, B.A. Aksoy, Y. Senbabaoglu, N. Schultz, C. Sander, Emerging landscape of oncogenic signatures across human cancers., Nat. Genet. 45 (2013) 1127–1133.

[7] P.D. Stenson, M. Mort, E. V Ball, K. Howells, A.D. Phillips, N.S. Thomas, D.N. Cooper, The Human Gene Mutation Database: 2008 update., Genome Med. 1 (2009) 13.

[8] P.J. Stephens, C.D. Greenman, B. Fu, F. Yang, G.R. Bignell, L.J. Mudie, E.D. Pleasance, K.W. Lau, D. Beare, L.A. Stebbings, S. McLaren, M.-L. Lin, D.J. McBride, I. Varela, S. Nik-Zainal, C. Leroy, M. Jia, A. Menzies, A.P. Butler, J.W. Teague, M.A. Quail, J. Burton, H. Swerdlow, N.P. Carter, L.A. Morsberger, C. Iacobuzio-Donahue, G.A. Follows, A.R. Green, A.M. Flanagan, M.R. Stratton, P.A. Futreal, P.J. Campbell, Massive genomic rearrangement acquired in a single catastrophic event during cancer development., Cell. 144 (2011) 27–40.

[9] D.L. Nicolae, E. Gamazon, W. Zhang, S. Duan, M.E. Dolan, N.J. Cox, Trait-associated SNPs are more likely to be eQTLs: annotation to enhance discovery from GWAS., PLoS Genet. 6 (2010) e1000888.

[10] A. Gusev, S.H. Lee, G. Trynka, H. Finucane, B.J. Vilhjálmsson, H. Xu, C. Zang, S. Ripke, B. Bulik-Sullivan, E. Stahl, Schizophrenia Working Group of the Psychiatric Genomics Consortium, SWE-SCZ Consortium, A.K. Kähler, C.M. Hultman, S.M. Purcell, S.A. McCarroll, M. Daly, B. Pasaniuc, P.F. Sullivan, B.M. Neale, N.R. Wray, S. Raychaudhuri, A.L. Price, Schizophrenia Working Group of the Psychiatric Genomics Consortium, SWE-SCZ Consortium, Partitioning heritability of regulatory and cell-type-specific variants across 11 common diseases., Am. J. Hum. Genet. 95 (2014) 535–552.

[11] M.T. Maurano, R. Humbert, E. Rynes, R.E. Thurman, E. Haugen, H. Wang, A.P. Reynolds, R. Sandstrom, H. Qu, J. Brody, A. Shafer, F. Neri, K. Lee, T. Kutyavin, S. Stehling-Sun, A.K. Johnson, T.K. Canfield, E. Giste, M. Diegel, D. Bates, R.S. Hansen, S. Neph, P.J. Sabo, S. Heimfeld, A. Raubitschek, S. Ziegler, C. Cotsapas, N. Sotoodehnia, I. Glass, S.R. Sunyaev, R. Kaul, J.A. Stamatoyannopoulos, Systematic localization of common disease-associated variation in regulatory DNA., Science. 337 (2012) 1190–1195.

[12] F.W. Huang, E. Hodis, M.J. Xu, G. V. Kryukov, L. Chin, L.A. Garraway, Highly recurrent TERT promoter mutations in human melanoma., Science. 339 (2013) 957–959.

[13] S. Horn, A. Figl, P.S. Rachakonda, C. Fischer, A. Sucker, A. Gast, S. Kadel, I. Moll, E. Nagore, K. Hemminki, D. Schadendorf, R. Kumar, TERT promoter mutations in familial and sporadic melanoma., Science. 339 (2013) 959–961.

[14] P.D. Stenson, E. V. Ball, M. Mort, A.D. Phillips, J.A. Shiel, N.S.T. Thomas, S. Abeysinghe, M. Krawczak, D.N. Cooper, Human Gene Mutation Database (HGMD): 2003 update, Hum. Mutat. 21 (2003) 577–581.

[15] S. Richards, N. Aziz, S. Bale, D. Bick, S. Das, J. Gastier-Foster, W.W. Grody, M. Hegde, E. Lyon, E. Spector, K. Voelkerding, H.L. Rehm, ACMG Laboratory Quality Assurance Committee, Standards and guidelines for the interpretation of sequence variants: a joint consensus recommendation of the American College of Medical Genetics and Genomics and the Association for Molecular Pathology., Genet. Med. 17 (2015) 405–423.

[16] G.M. Cooper, J. Shendure, Needles in stacks of needles: finding disease-causal variants in a wealth of genomic data., Nat. Rev. Genet. 12 (2011) 628–640.

[17] P. Yue, J. Moult, Identification and analysis of deleterious human SNPs., J. Mol. Biol. 356 (2006) 1263–1274.

[18] C. Douville, D.L. Masica, P.D. Stenson, D.N. Cooper, D.M. Gygax, R. Kim, M. Ryan, R. Karchin, Assessing the Pathogenicity of Insertion and Deletion Variants with the Variant Effect Scoring Tool (VEST-Indel)., Hum. Mutat. 37 (2016) 28–35.

[19] M. Kircher, D. Witten, P. Jain, B. O’Roak, G. Cooper, J. Shendure, A general framework for estimating the relative pathogenicity of human genetic variants., Nat. Genet. 46 (2014) 310–315.

[20] I.A. Adzhubei, S. Schmidt, L. Peshkin, V.E. Ramensky, A. Gerasimova, P. Bork, A.S. Kondrashov, S.R. Sunyaev, A method and server for predicting damaging missense mutations., Nat. Methods. 7 (2010) 248–249.

[21] H. Carter, C. Douville, P.D. Stenson, D.N. Cooper, R. Karchin, Identifying Mendelian disease genes with the variant effect scoring tool., BMC Genomics. 14 Suppl 3 (2013) S3.

[22] O. Lichtarge, H.R. Bourne, F.E. Cohen, An Evolutionary Trace Method Defines Binding Surfaces Common to Protein Families., J. Mol. Biol. 257 (1996) 342–358.

[23] P.C. Ng, S. Henikoff, SIFT: Predicting amino acid changes that affect protein function., Nucleic Acids Res. 31 (2003) 3812–3814.

[24] Z. Wang, J. Moult, SNPs, protein structure, and disease., Hum. Mutat. 17 (2001) 263–270.

[25] P. Yue, Z. Li, J. Moult, Loss of protein structure stability as a major causative factor in monogenic disease., J. Mol. Biol. 353 (2005) 459–473.

[26] Z. Shi, J. Moult, Structural and Functional Impact of Cancer-Related Missense Somatic Mutations., J. Mol. Biol. 413 (2011) 495–512.

[27] Y. Xue, Y. Chen, Q. Ayub, N. Huang, E. V. Ball, M. Mort, A.D. Phillips, K. Shaw, P.D. Stenson, D.N. Cooper, C. Tyler-Smith, 1000 Genomes Project Consortium, Deleterious- and disease-allele prevalence in healthy individuals: insights from current predictions, mutation databases, and population-scale resequencing., Am. J. Hum. Genet. 91 (2012) 1022–1032.

[28] L.B. Alexandrov, S. Nik-Zainal, D.C. Wedge, S.A.J.R. Aparicio, S. Behjati, A. V. Biankin, G.R. Bignell, N. Bolli, A. Borg, A.-L. Børresen-Dale, S. Boyault, B. Burkhardt, A.P. Butler, C. Caldas, H.R. Davies, C. Desmedt, R. Eils, J.E. Eyfjörd, J.A. Foekens, M. Greaves, F. Hosoda, B. Hutter, T. Ilicic, S. Imbeaud, M. Imielinsk, N. Jäger, D.T.W. Jones, D. Jones, S. Knappskog, M. Kool, S.R. Lakhani, C. López-Otín, S. Martin, N.C. Munshi, H. Nakamura, P.A. Northcott, M. Pajic, E. Papaemmanuil, A. Paradiso, J. V. Pearson, X.S. Puente, K. Raine, M. Ramakrishna, A.L. Richardson, J. Richter, P. Rosenstiel, M. Schlesner, T.N. Schumacher, P.N. Span, J.W. Teague, Y. Totoki, A.N.J. Tutt, R. Valdés-Mas, M.M. van Buuren, L. van ’t Veer, A. Vincent-Salomon, N. Waddell, L.R. Yates, J. Zucman-Rossi, P. Andrew Futreal, U. McDermott, P. Lichter, M. Meyerson, S.M. Grimmond, R. Siebert, E. Campo, T. Shibata, S.M. Pfister, P.J. Campbell, M.R. Stratton, E. Campo, T. Shibata, S.M. Pfister, P.J. Campbell, M.R. Stratton, Signatures of mutational processes in human cancer, Nature. 500 (2013) 415–421. doi:10.1038/nature12477.

[29] M.S. Lawrence, P. Stojanov, C.H. Mermel, J.T. Robinson, L.A. Garraway, T.R. Golub, M. Meyerson, S.B. Gabriel, E.S. Lander, G. Getz, Discovery and saturation analysis of cancer genes across 21 tumour types., Nature. 505 (2014) 495–501.

[30] P. Armitage, R. Doll, The age distribution of cancer and a multi-stage theory of carcinogenesis., Br. J. Cancer. 8 (1954) 1–12.

[31] R. Sabarinathan, O. Pich, I. Martincorena, C. Rubio-Perez, M. Juul, J. Wala, S. Schumacher, O. Shapira, N. Sidiropoulos, S. Waszak, D. Tamborero, L. Mularoni, E. Rheinbay, H. Hornshoj, J. Deu-Pons, F. Muinos, J. Bertl, Q. Guo, J. Weischenfeldt, J.O. Korbel, G. Getz, P.J. Campbell, J.S. Pedersen, R. Beroukhim, A. Perez-Gonzalez, N. Lopez-Bigas, PCAWG Drivers and Functional Interpretation Group, I.P.-C.A. of W.G. Net, The whole-genome panorama of cancer drivers., bioRxiv. (2017) 190330.

[32] D. Tamborero, A. Gonzalez-Perez, N. Lopez-Bigas, OncodriveCLUST: exploiting the positional clustering of somatic mutations to identify cancer genes., Bioinformatics. 29 (2013) 2238–2244.

[33] D. Tamborero, A. Gonzalez-Perez, C. Perez-Llamas, J. Deu-Pons, C. Kandoth, J. Reimand, M.S. Lawrence, G. Getz, G.D. Bader, L. Ding, N. Lopez-Bigas, Comprehensive identification of mutational cancer driver genes across 12 tumor types., Sci. Rep. 3 (2013) 2650.

[34] J. Reimand, G.D. Bader, Systematic analysis of somatic mutations in phosphorylation signaling predicts novel cancer drivers., Mol. Syst. Biol. 9 (2013) 637.

[35] N.D. Dees, Q. Zhang, C. Kandoth, M.C. Wendl, W. Schierding, D.C. Koboldt, T.B. Mooney, M.B. Callaway, D. Dooling, E.R. Mardis, R.K. Wilson, L. Ding, MuSiC: identifying mutational significance in cancer genomes., Genome Res. 22 (2012) 1589–1598.

[36] A. Gonzalez-Perez, N. Lopez-Bigas, Functional impact bias reveals cancer drivers., Nucleic Acids Res. 40 (2012) e169.

[37] C. Rubio-Perez, D. Tamborero, M.P. Schroeder, A.A. Antolín, J. Deu-Pons, C. Perez-Llamas, J. Mestres, A. Gonzalez-Perez, N. Lopez-Bigas, In silico prescription of anticancer drugs to cohorts of 28 tumor types reveals targeting opportunities., Cancer Cell. 27 (2015) 382–396.

[38] M.D.M. Leiserson, F. Vandin, H.-T. Wu, J.R. Dobson, J. V Eldridge, J.L. Thomas, A. Papoutsaki, Y. Kim, B. Niu, M. McLellan, M.S. Lawrence, A. Gonzalez-Perez, D. Tamborero, Y. Cheng, G.A. Ryslik, N. Lopez-Bigas, G. Getz, L. Ding, B.J. Raphael, Pan-cancer network analysis identifies combinations of rare somatic mutations across pathways and protein complexes., Nat. Genet. 47 (2015) 106–114.

[39] T. Davoli, A.W. Xu, K.E. Mengwasser, L.M. Sack, J.C. Yoon, P.J. Park, S.J. Elledge, Cumulative haploinsufficiency and triplosensitivity drive aneuploidy patterns and shape the cancer genome., Cell. 155 (2013) 948–962.

[40] C.H. Mermel, S.E. Schumacher, B. Hill, M.L. Meyerson, R. Beroukhim, G. Getz, GISTIC2.0 facilitates sensitive and confident localization of the targets of focal somatic copy-number alteration in human cancers., Genome Biol. 12 (2011) R41.

[41] C.J. Tokheim, N. Papadopoulos, K.W. Kinzler, B. Vogelstein, R. Karchin, Evaluating the evaluation of cancer driver genes., Proc. Natl. Acad. Sci. U. S. A. 113 (2016) 14330–14335.

[42] P.A. Futreal, L. Coin, M. Marshall, T. Down, T. Hubbard, R. Wooster, N. Rahman, M.R. Stratton, A census of human cancer genes., Nat. Rev. Cancer. 4 (2004) 177–183.

[43] H. Carter, S. Chen, L. Isik, S. Tyekucheva, V.E. Velculescu, K.W. Kinzler, B. Vogelstein, R. Karchin, Cancer-specific high-throughput annotation of somatic mutations: computational prediction of driver missense mutations., Cancer Res. 69 (2009) 6660–6667.

[44] B. Reva, Y. Antipin, C. Sander, Predicting the functional impact of protein mutations: application to cancer genomics., Nucleic Acids Res. 39 (2011) e118.

[45] Y. Mao, H. Chen, H. Liang, F. Meric-Bernstam, G.B. Mills, K. Chen, CanDrA: cancer-specific driver missense mutation annotation with optimized features., PLoS One. 8 (2013) e77945.

[46] J.S. Kaminker, Y. Zhang, A. Waugh, P.M. Haverty, B. Peters, D. Sebisanovic, J. Stinson, W.F. Forrest, J.F. Bazan, S. Seshagiri, Z. Zhang, Distinguishing cancer-associated missense mutations from common polymorphisms., Cancer Res. 67 (2007) 465–473.

[47] A. Gonzalez-Perez, J. Deu-Pons, N. Lopez-Bigas, Improving the prediction of the functional impact of cancer mutations by baseline tolerance transformation., Genome Med. 4 (2012) 89.

[48] H.A. Shihab, J. Gough, D.N. Cooper, I.N.M. Day, T.R. Gaunt, Predicting the functional consequences of cancer-associated amino acid substitutions., Bioinformatics. 29 (2013) 1504–1510.

[49] D. Tamborero, C. Rubio-Perez, J. Deu-Pons, M.P. Schroeder, A. Vivancos, A. Rovira, I. Tusquets, J. Albanell, J. Rodon, J. Tabernero, C. de Torres, R. Dienstmann, A. Gonzalez-Perez, N. Lopez-Bigas, Cancer Genome Interpreter annotates the biological and clinical relevance of tumor alterations, Genome Med. 10 (2018) 25.

[50] V. Vacic, P.R.L. Markwick, C.J. Oldfield, X. Zhao, C. Haynes, V.N. Uversky, L.M. Iakoucheva, Disease-associated mutations disrupt functionally important regions of intrinsic protein disorder., PLoS Comput. Biol. 8 (2012) e1002709.

[51] U. Midic, C.J. Oldfield, A.K. Dunker, Z. Obradovic, V.N. Uversky, Protein disorder in the human diseasome: unfoldomics of human genetic diseases., BMC Genomics. 10 Suppl 1 (2009) S12.

[52] Cancer Genome Atlas Network, D.C. Koboldt, R.S. Fulton, M.D. McLellan, H. Schmidt, J. Kalicki-Veizer, J.F. McMichael, L.L. Fulton, D.J. Dooling, L. Ding, E.R. Mardis, R.K. Wilson, A. Ally, M. Balasundaram, Y.S.N. Butterfield, R. Carlsen, C. Carter, A. Chu, E. Chuah, H.-J.E. Chun, R.J.N. Coope, N. Dhalla, R. Guin, C. Hirst, M. Hirst, R.A. Holt, D. Lee, H.I. Li, M. Mayo, R.A. Moore, A.J. Mungall, E. Pleasance, A. Gordon Robertson, J.E. Schein, A. Shafiei, P. Sipahimalani, J.R. Slobodan, D. Stoll, A. Tam, N. Thiessen, R.J. Varhol, N. Wye, T. Zeng, Y. Zhao, I. Birol, S.J.M. Jones, M.A. Marra, A.D. Cherniack, G. Saksena, R.C. Onofrio, N.H. Pho, S.L. Carter, S.E. Schumacher, B. Tabak, B. Hernandez, J. Gentry, H. Nguyen, A. Crenshaw, K. Ardlie, R. Beroukhim, W. Winckler, G. Getz, S.B. Gabriel, M. Meyerson, L. Chin, P.J. Park, R. Kucherlapati, K.A. Hoadley, J. Todd Auman, C. Fan, Y.J. Turman, Y. Shi, L. Li, M.D. Topal, X. He, H.-H. Chao, A. Prat, G.O. Silva, M.D. Iglesia, W. Zhao, J. Usary, J.S. Berg, M. Adams, J. Booker, J. Wu, A. Gulabani, T. Bodenheimer, A.P. Hoyle, J. V. Simons, M.G. Soloway, L.E. Mose, S.R. Jefferys, S. Balu, J.S. Parker, D. Neil Hayes, C.M. Perou, S. Malik, S. Mahurkar, H. Shen, D.J. Weisenberger, T. Triche Jr, P.H. Lai, M.S. Bootwalla, D.T. Maglinte, B.P. Berman, D.J. Van Den Berg, S.B. Baylin, P.W. Laird, C.J. Creighton, L.A. Donehower, G. Getz, M. Noble, D. Voet, G. Saksena, N. Gehlenborg, D. DiCara, J. Zhang, H. Zhang, C.-J. Wu, S. Yingchun Liu, M.S. Lawrence, L. Zou, A. Sivachenko, P. Lin, P. Stojanov, R. Jing, J. Cho, R. Sinha, R.W. Park, M.-D. Nazaire, J. Robinson, H. Thorvaldsdottir, J. Mesirov, P.J. Park, L. Chin, S. Reynolds, R.B. Kreisberg, B. Bernard, R. Bressler, T. Erkkila, J. Lin, V. Thorsson, W. Zhang, I. Shmulevich, G. Ciriello, N. Weinhold, N. Schultz, J. Gao, E. Cerami, B. Gross, A. Jacobsen, R. Sinha, B. Arman Aksoy, Y. Antipin, B. Reva, R. Shen, B.S. Taylor, M. Ladanyi, C. Sander, P. Anur, P.T. Spellman, Y. Lu, W. Liu, R.R.G. Verhaak, G.B. Mills, R. Akbani, N. Zhang, B.M. Broom, T.D. Casasent, C. Wakefield, A.K. Unruh, K. Baggerly, K. Coombes, J.N. Weinstein, D. Haussler, C.C. Benz, J.M. Stuart, S.C. Benz, J. Zhu, C.C. Szeto, G.K. Scott, C. Yau, E.O. Paull, D. Carlin, C. Wong, A. Sokolov, J. Thusberg, S. Mooney, S. Ng, T.C. Goldstein, K. Ellrott, M. Grifford, C. Wilks, S. Ma, B. Craft, C. Yan, Y. Hu, D. Meerzaman, J.M. Gastier-Foster, J. Bowen, N.C. Ramirez, A.D. Black, R.E. XPATH ERROR: unknown variable “tname”., P. White, E.J. Zmuda, J. Frick, T.M. Lichtenberg, R. Brookens, M.M. George, M.A. Gerken, H.A. Harper, K.M. Leraas, L.J. Wise, T.R. Tabler, C. McAllister, T. Barr, M. Hart-Kothari, K. Tarvin, C. Saller, G. Sandusky, C. Mitchell, M. V. Iacocca, J. Brown, B. Rabeno, C. Czerwinski, N. Petrelli, O. Dolzhansky, M. Abramov, O. Voronina, O. Potapova, J.R. Marks, W.M. Suchorska, D. Murawa, W. Kycler, M. Ibbs, K. Korski, A. Spychała, P. Murawa, J.J. Brzeziński, H. Perz, R. Łaźniak, M. Teresiak, H. Tatka, E. Leporowska, M. Bogusz-Czerniewicz, J. Malicki, A. Mackiewicz, M. Wiznerowicz, X. Van Le, B. Kohl, N. Viet Tien, R. Thorp, N. Van Bang, H. Sussman, B. Duc Phu, R. Hajek, N. Phi Hung, T. Viet The Phuong, H. Quyet Thang, K. Zaki Khan, R. Penny, D. Mallery, E. Curley, C. Shelton, P. Yena, J.N. Ingle, F.J. Couch, W.L. Lingle, T.A. King, A. Maria Gonzalez-Angulo, G.B. Mills, M.D. Dyer, S. Liu, X. Meng, M. Patangan, F. Waldman, H. Stöppler, W. Kimryn Rathmell, L. Thorne, M. Huang, L. Boice, A. Hill, C. Morrison, C. Gaudioso, W. Bshara, K. Daily, S.C. Egea, M.D. Pegram, C. Gomez-Fernandez, R. Dhir, R. Bhargava, A. Brufsky, C.D. Shriver, J.A. Hooke, J. Leigh Campbell, R.J. Mural, H. Hu, S. Somiari, C. Larson, B. Deyarmin, L. Kvecher, A.J. Kovatich, M.J. Ellis, T.A. King, H. Hu, F.J. Couch, R.J. Mural, T. Stricker, K. White, O. Olopade, J.N. Ingle, C. Luo, Y. Chen, J.R. Marks, F. Waldman, M. Wiznerowicz, R. Bose, L.-W. Chang, A.H. Beck, A. Maria Gonzalez-Angulo, T. Pihl, M. Jensen, R. Sfeir, A. Kahn, A. Chu, P. Kothiyal, Z. Wang, E. Snyder, J. Pontius, B. Ayala, M. Backus, J. Walton, J. Baboud, D. Berton, M. Nicholls, D. Srinivasan, R. Raman, S. Girshik, P. Kigonya, S. Alonso, R. Sanbhadti, S. Barletta, D. Pot, M. Sheth, J.A. Demchok, K.R. Mills Shaw, L. Yang, G. Eley, M.L. Ferguson, R.W. Tarnuzzer, J. Zhang, L.A.L. Dillon, K. Buetow, P. Fielding, B.A. Ozenberger, M.S. Guyer, H.J. Sofia, J.D. Palchik, Comprehensive molecular portraits of human breast tumours., Nature. 490 (2012) 61–70.

[53] Cancer Genome Atlas Research Network, R. McLendon, A. Friedman, D. Bigner, E.G. Van Meir, D.J. Brat, G. M. Mastrogianakis, J.J. Olson, T. Mikkelsen, N. Lehman, K. Aldape, W.K. Alfred Yung, O. Bogler, S. VandenBerg, M. Berger, M. Prados, D. Muzny, M. Morgan, S. Scherer, A. Sabo, L. Nazareth, L. Lewis, O. Hall, Y. Zhu, Y. Ren, O. Alvi, J. Yao, A. Hawes, S. Jhangiani, G. Fowler, A. San Lucas, C. Kovar, A. Cree, H. Dinh, J. Santibanez, V. Joshi, M.L. Gonzalez-Garay, C.A. Miller, A. Milosavljevic, L. Donehower, D.A. Wheeler, R.A. Gibbs, K. Cibulskis, C. Sougnez, T. Fennell, S. Mahan, J. Wilkinson, L. Ziaugra, R. Onofrio, T. Bloom, R. Nicol, K. Ardlie, J. Baldwin, S. Gabriel, E.S. Lander, L. Ding, R.S. Fulton, M.D. McLellan, J. Wallis, D.E. Larson, X. Shi, R. Abbott, L. Fulton, K. Chen, D.C. Koboldt, M.C. Wendl, R. Meyer, Y. Tang, L. Lin, J.R. Osborne, B.H. Dunford-Shore, T.L. Miner, K. Delehaunty, C. Markovic, G. Swift, W. Courtney, C. Pohl, S. Abbott, A. Hawkins, S. Leong, C. Haipek, H. Schmidt, M. Wiechert, T. Vickery, S. Scott, D.J. Dooling, A. Chinwalla, G.M. Weinstock, E.R. Mardis, R.K. Wilson, G. Getz, W. Winckler, R.G.W. Verhaak, M.S. Lawrence, M. O’Kelly, J. Robinson, G. Alexe, R. Beroukhim, S. Carter, D. Chiang, J. Gould, S. Gupta, J. Korn, C. Mermel, J. Mesirov, S. Monti, H. Nguyen, M. Parkin, M. Reich, N. Stransky, B.A. Weir, L. Garraway, T. Golub, M. Meyerson, L. Chin, A. Protopopov, J. Zhang, I. Perna, S. Aronson, N. Sathiamoorthy, G. Ren, J. Yao, W.R. Wiedemeyer, H. Kim, S. Won Kong, Y. Xiao, I.S. Kohane, J. Seidman, P.J. Park, R. Kucherlapati, P.W. Laird, L. Cope, J.G. Herman, D.J. Weisenberger, F. Pan, D. Van Den Berg, L. Van Neste, J. Mi Yi, K.E. Schuebel, S.B. Baylin, D.M. Absher, J.Z. Li, A. Southwick, S. Brady, A. Aggarwal, T. Chung, G. Sherlock, J.D. Brooks, R.M. Myers, P.T. Spellman, E. Purdom, L.R. Jakkula, A. V. Lapuk, H. Marr, S. Dorton, Y. Gi Choi, J. Han, A. Ray, V. Wang, S. Durinck, M. Robinson, N.J. Wang, K. Vranizan, V. Peng, E. Van Name, G. V. Fontenay, J. Ngai, J.G. Conboy, B. Parvin, H.S. Feiler, T.P. Speed, J.W. Gray, C. Brennan, N.D. Socci, A. Olshen, B.S. Taylor, A. Lash, N. Schultz, B. Reva, Y. Antipin, A. Stukalov, B. Gross, E. Cerami, W. Qing Wang, L.-X. Qin, V.E. Seshan, L. Villafania, M. Cavatore, L. Borsu, A. Viale, W. Gerald, C. Sander, M. Ladanyi, C.M. Perou, D. Neil Hayes, M.D. Topal, K.A. Hoadley, Y. Qi, S. Balu, Y. Shi, J. Wu, R. Penny, M. Bittner, T. Shelton, E. Lenkiewicz, S. Morris, D. Beasley, S. Sanders, A. Kahn, R. Sfeir, J. Chen, D. Nassau, L. Feng, E. Hickey, J. Zhang, J.N. Weinstein, A. Barker, D.S. Gerhard, J. Vockley, C. Compton, J. Vaught, P. Fielding, M.L. Ferguson, C. Schaefer, S. Madhavan, K.H. Buetow, F. Collins, P. Good, M. Guyer, B. Ozenberger, J. Peterson, E. Thomson, Comprehensive genomic characterization defines human glioblastoma genes and core pathways., Nature. 455 (2008) 1061–1068.

[54] Cancer Genome Atlas Research Network, D. Bell, A. Berchuck, M. Birrer, J. Chien, D.W. Cramer, F. Dao, R. Dhir, P. DiSaia, H. Gabra, P. Glenn, A.K. Godwin, J. Gross, L. Hartmann, M. Huang, D.G. Huntsman, M. Iacocca, M. Imielinski, S. Kalloger, B.Y. Karlan, D.A. Levine, G.B. Mills, C. Morrison, D. Mutch, N. Olvera, S. Orsulic, K. Park, N. Petrelli, B. Rabeno, J.S. Rader, B.I. Sikic, K. Smith-McCune, A.K. Sood, D. Bowtell, R. Penny, J.R. Testa, K. Chang, H.H. Dinh, J.A. Drummond, G. Fowler, P. Gunaratne, A.C. Hawes, C.L. Kovar, L.R. Lewis, M.B. Morgan, I.F. Newsham, J. Santibanez, J.G. Reid, L.R. Trevino, Y.-Q. Wu, M. Wang, D.M. Muzny, D.A. Wheeler, R.A. Gibbs, G. Getz, M.S. Lawrence, K. Cibulskis, A.Y. Sivachenko, C. Sougnez, D. Voet, J. Wilkinson, T. Bloom, K. Ardlie, T. Fennell, J. Baldwin, S. Gabriel, E.S. Lander, L. Ding, R.S. Fulton, D.C. Koboldt, M.D. McLellan, T. Wylie, J. Walker, M. O’Laughlin, D.J. Dooling, L. Fulton, R. Abbott, N.D. Dees, Q. Zhang, C. Kandoth, M. Wendl, W. Schierding, D. Shen, C.C. Harris, H. Schmidt, J. Kalicki, K.D. Delehaunty, C.C. Fronick, R. Demeter, L. Cook, J.W. Wallis, L. Lin, V.J. Magrini, J.S. Hodges, J.M. Eldred, S.M. Smith, C.S. Pohl, F. Vandin, B.J. Raphael, G.M. Weinstock, E.R. Mardis, R.K. Wilson, M. Meyerson, W. Winckler, G. Getz, R.G.W. Verhaak, S.L. Carter, C.H. Mermel, G. Saksena, H. Nguyen, R.C. Onofrio, M.S. Lawrence, D. Hubbard, S. Gupta, A. Crenshaw, A.H. Ramos, K. Ardlie, L. Chin, A. Protopopov, J. Zhang, T.M. Kim, I. Perna, Y. Xiao, H. Zhang, G. Ren, N. Sathiamoorthy, R.W. Park, E. Lee, P.J. Park, R. Kucherlapati, D.M. Absher, L. Waite, G. Sherlock, J.D. Brooks, J.Z. Li, J. Xu, R.M. Myers, P.W. Laird, L. Cope, J.G. Herman, H. Shen, D.J. Weisenberger, H. Noushmehr, F. Pan, T. Triche Jr, B.P. Berman, D.J. Van Den Berg, J. Buckley, S.B. Baylin, P.T. Spellman, E. Purdom, P. Neuvial, H. Bengtsson, L.R. Jakkula, S. Durinck, J. Han, S. Dorton, H. Marr, Y.G. Choi, V. Wang, N.J. Wang, J. Ngai, J.G. Conboy, B. Parvin, H.S. Feiler, T.P. Speed, J.W. Gray, D.A. Levine, N.D. Socci, Y. Liang, B.S. Taylor, N. Schultz, L. Borsu, A.E. Lash, C. Brennan, A. Viale, C. Sander, M. Ladanyi, K.A. Hoadley, S. Meng, Y. Du, Y. Shi, L. Li, Y.J. Turman, D. Zang, E.B. Helms, S. Balu, X. Zhou, J. Wu, M.D. Topal, D.N. Hayes, C.M. Perou, G. Getz, D. Voet, G. Saksena, J. Zhang, H. Zhang, C.J. Wu, S. Shukla, K. Cibulskis, M.S. Lawrence, A. Sivachenko, R. Jing, R.W. Park, Y. Liu, P.J. Park, M. Noble, L. Chin, H. Carter, D. Kim, R. Karchin, P.T. Spellman, E. Purdom, P. Neuvial, H. Bengtsson, S. Durinck, J. Han, J.E. Korkola, L.M. Heiser, R.J. Cho, Z. Hu, B. Parvin, T.P. Speed, J.W. Gray, N. Schultz, E. Cerami, B.S. Taylor, A. Olshen, B. Reva, Y. Antipin, R. Shen, P. Mankoo, R. Sheridan, G. Ciriello, W.K. Chang, J.A. Bernanke, L. Borsu, D.A. Levine, M. Ladanyi, C. Sander, D. Haussler, C.C. Benz, J.M. Stuart, S.C. Benz, J.Z. Sanborn, C.J. Vaske, J. Zhu, C. Szeto, G.K. Scott, C. Yau, K.A. Hoadley, Y. Du, S. Balu, D.N. Hayes, C.M. Perou, M.D. Wilkerson, N. Zhang, R. Akbani, K.A. Baggerly, W.K. Yung, G.B. Mills, J.N. Weinstein, R. Penny, T. Shelton, D. Grimm, M. Hatfield, S. Morris, P. Yena, P. Rhodes, M. Sherman, J. Paulauskis, S. Millis, A. Kahn, J.M. Greene, R. Sfeir, M.A. Jensen, J. Chen, J. Whitmore, S. Alonso, J. Jordan, A. Chu, J. Zhang, A. Barker, C. Compton, G. Eley, M. Ferguson, P. Fielding, D.S. Gerhard, R. Myles, C. Schaefer, K.R. Mills Shaw, J. Vaught, J.B. Vockley, P.J. Good, M.S. Guyer, B. Ozenberger, J. Peterson, E. Thomson, Integrated genomic analyses of ovarian carcinoma., Nature. 474 (2011) 609–615.

[55] Cancer Genome Atlas Research Network, G. Getz, S.B. Gabriel, K. Cibulskis, E. Lander, A. Sivachenko, C. Sougnez, M. Lawrence, C. Kandoth, D. Dooling, R. Fulton, L. Fulton, J. Kalicki-Veizer, M.D. McLellan, M. O’Laughlin, H. Schmidt, R.K. Wilson, K. Ye, L. Ding, E.R. Mardis, A. Ally, M. Balasundaram, I. Birol, Y.S.N. Butterfield, R. Carlsen, C. Carter, A. Chu, E. Chuah, H.-J.E. Chun, N. Dhalla, R. Guin, C. Hirst, R.A. Holt, S.J.M. Jones, D. Lee, H.I. Li, M.A. Marra, M. Mayo, R.A. Moore, A.J. Mungall, P. Plettner, J.E. Schein, P. Sipahimalani, A. Tam, R.J. Varhol, A. Gordon Robertson, A.D. Cherniack, I. Pashtan, G. Saksena, R.C. Onofrio, S.E. Schumacher, B. Tabak, S.L. Carter, B. Hernandez, J. Gentry, H.B. Salvesen, K. Ardlie, G. Getz, W. Winckler, R. Beroukhim, S.B. Gabriel, M. Meyerson, A. Hadjipanayis, S. Lee, H.S. Mahadeshwar, P. Park, A. Protopopov, X. Ren, S. Seth, X. Song, J. Tang, R. Xi, L. Yang, D. Zeng, R. Kucherlapati, L. Chin, J. Zhang, J. Todd Auman, S. Balu, T. Bodenheimer, E. Buda, D. Neil Hayes, A.P. Hoyle, S.R. Jefferys, C.D. Jones, S. Meng, P.A. Mieczkowski, L.E. Mose, J.S. Parker, C.M. Perou, J. Roach, Y. Shi, J. V. Simons, M.G. Soloway, D. Tan, M.D. Topal, S. Waring, J. Wu, K.A. Hoadley, S.B. Baylin, M.S. Bootwalla, P.H. Lai, T.J. Triche Jr, D.J. Van Den Berg, D.J. Weisenberger, P.W. Laird, H. Shen, L. Chin, J. Zhang, G. Getz, J. Cho, D. DiCara, S. Frazer, D. Heiman, R. Jing, P. Lin, W. Mallard, P. Stojanov, D. Voet, H. Zhang, L. Zou, M. Noble, M. Lawrence, S.M. Reynolds, I. Shmulevich, B. Arman Aksoy, Y. Antipin, G. Ciriello, G. Dresdner, J. Gao, B. Gross, A. Jacobsen, M. Ladanyi, B. Reva, C. Sander, R. Sinha, S. Onur Sumer, B.S. Taylor, E. Cerami, N. Weinhold, N. Schultz, R. Shen, S. Benz, T. Goldstein, D. Haussler, S. Ng, C. Szeto, J. Stuart, C.C. Benz, C. Yau, W. Zhang, M. Annala, B.M. Broom, T.D. Casasent, Z. Ju, H. Liang, G. Liu, Y. Lu, A.K. Unruh, C. Wakefield, J.N. Weinstein, N. Zhang, Y. Liu, R. Broaddus, R. Akbani, G.B. Mills, C. Adams, T. Barr, A.D. Black, J. Bowen, J. Deardurff, J. Frick, J.M. Gastier-Foster, T. Grossman, H.A. Harper, M. Hart-Kothari, C. Helsel, A. Hobensack, H. Kuck, K. Kneile, K.M. Leraas, T.M. Lichtenberg, C. McAllister, R.E. Pyatt, N.C. Ramirez, T.R. Tabler, N. Vanhoose, P. White, L. Wise, E. Zmuda, N. Barnabas, C. Berry-Green, V. Blanc, L. Boice, M. Button, A. Farkas, A. Green, J. MacKenzie, D. Nicholson, S.E. Kalloger, C. Blake Gilks, B.Y. Karlan, J. Lester, S. Orsulic, M. Borowsky, M. Cadungog, C. Czerwinski, L. Huelsenbeck-Dill, M. Iacocca, N. Petrelli, B. Rabeno, G. Witkin, E. Nemirovich-Danchenko, O. Potapova, D. Rotin, A. Berchuck, M. Birrer, P. DiSaia, L. Monovich, E. Curley, J. Gardner, D. Mallery, R. Penny, S.C. Dowdy, B. Winterhoff, L. Dao, B. Gostout, A. Meuter, A. Teoman, F. Dao, N. Olvera, F. Bogomolniy, K. Garg, R.A. Soslow, D.A. Levine, M. Abramov, J.M.S. Bartlett, S. Kodeeswaran, J. Parfitt, F. Moiseenko, B.A. Clarke, M.T. Goodman, M.E. Carney, R.K. Matsuno, J. Fisher, M. Huang, W. Kimryn Rathmell, L. Thorne, L. Van Le, R. Dhir, R. Edwards, E. Elishaev, K. Zorn, R. Broaddus, P.J. Goodfellow, D. Mutch, N. Schultz, Y. Liu, R. Akbani, A.D. Cherniack, E. Cerami, N. Weinhold, H. Shen, K.A. Hoadley, A.B. Kahn, D.W. Bell, P.M. Pollock, C. Wang, D. A. Wheeler, E. Shinbrot, B.Y. Karlan, A. Berchuck, S.C. Dowdy, B. Winterhoff, M.T. Goodman, A. Gordon Robertson, R. Beroukhim, I. Pashtan, H.B. Salvesen, P.W. Laird, M. Noble, J. Stuart, L. Ding, C. Kandoth, C. Blake Gilks, R.A. Soslow, P.J. Goodfellow, D. Mutch, R. Broaddus, W. Zhang, G.B. Mills, R. Kucherlapati, E.R. Mardis, D.A. Levine, B. Ayala, A.L. Chu, M.A. Jensen, P. Kothiyal, T.D. Pihl, J. Pontius, D.A. Pot, E.E. Snyder, D. Srinivasan, A.B. Kahn, K.R. Mills Shaw, M. Sheth, T. Davidsen, G. Eley Martin L. Ferguson, J.A. Demchok, L. Yang, M.S. Guyer, B.A. Ozenberger, H.J. Sofia, C. Kandoth, N. Schultz, A.D. Cherniack, R. Akbani, Y. Liu, H. Shen, A. Gordon Robertson, I. Pashtan, R. Shen, C.C. Benz, C. Yau, P.W. Laird, L. Ding, W. Zhang, G.B. Mills, R. Kucherlapati, E.R. Mardis, D.A. Levine, Integrated genomic characterization of endometrial carcinoma., Nature. 497 (2013) 67–73.

[56] Cancer Genome Atlas Research Network, T.J. Ley, C. Miller, L. Ding, B.J. Raphael, A.J. Mungall, A.G. Robertson, K. Hoadley, T.J. Triche, P.W. Laird, J.D. Baty, L.L. Fulton, R. Fulton, S.E. Heath, J. Kalicki-Veizer, C. Kandoth, J.M. Klco, D.C. Koboldt, K.-L. Kanchi, S. Kulkarni, T.L. Lamprecht, D.E. Larson, L. Lin, C. Lu, M.D. McLellan, J.F. McMichael, J. Payton, H. Schmidt, D.H. Spencer, M.H. Tomasson, J.W. Wallis, L.D. Wartman, M.A. Watson, J. Welch, M.C. Wendl, A. Ally, M. Balasundaram, I. Birol, Y. Butterfield, R. Chiu, A. Chu, E. Chuah, H.-J. Chun, R. Corbett, N. Dhalla, R. Guin, A. He, C. Hirst, M. Hirst, R.A. Holt, S. Jones, A. Karsan, D. Lee, H.I. Li, M.A. Marra, M. Mayo, R.A. Moore, K. Mungall, J. Parker, E. Pleasance, P. Plettner, J. Schein, D. Stoll, L. Swanson, A. Tam, N. Thiessen, R. Varhol, N. Wye, Y. Zhao, S. Gabriel, G. Getz, C. Sougnez, L. Zou, M.D.M. Leiserson, F. Vandin, H.-T. Wu, F. Applebaum, S.B. Baylin, R. Akbani, B.M. Broom, K. Chen, T.C. Motter, K. Nguyen, J.N. Weinstein, N. Zhang, M.L. Ferguson, C. Adams, A. Black, J. Bowen, J. Gastier-Foster, T. Grossman, T. Lichtenberg, L. Wise, T. Davidsen, J.A. Demchok, K.R.M. Shaw, M. Sheth, H.J. Sofia, L. Yang, J.R. Downing, G. Eley, Genomic and epigenomic landscapes of adult de novo acute myeloid leukemia., N. Engl. J. Med. 368 (2013) 2059–2074.

[57] Cancer Genome Atlas Network, M.N. Bainbridge, K. Chang, H.H. Dinh, J.A. Drummond, G. Fowler, C.L. Kovar, L.R. Lewis, M.B. Morgan, I.F. Newsham, J.G. Reid, J. Santibanez, E. Shinbrot, L.R. Trevino, Y.-Q. Wu, M. Wang, P. Gunaratne, L.A. Donehower, C.J. Creighton, D.A. Wheeler, R.A. Gibbs, M.S. Lawrence, D. Voet, R. Jing, K. Cibulskis, A. Sivachenko, P. Stojanov, A. McKenna, E.S. Lander, S. Gabriel, G. Getz, L. Ding, R.S. Fulton, D.C. Koboldt, T. Wylie, J. Walker, D.J. Dooling, L. Fulton, K.D. Delehaunty, C.C. Fronick, R. Demeter, E.R. Mardis, R.K. Wilson, A. Chu, H.-J.E. Chun, A.J. Mungall, E. Pleasance, A. Gordon Robertson, D. Stoll, M. Balasundaram, I. Birol, Y.S.N. Butterfield, E. Chuah, R.J.N. Coope, N. Dhalla, R. Guin, C. Hirst, M. Hirst, R.A. Holt, D. Lee, H.I. Li, M. Mayo, R.A. Moore, J.E. Schein, J.R. Slobodan, A. Tam, N. Thiessen, R. Varhol, T. Zeng, Y. Zhao, S.J.M. Jones, M.A. Marra, A.J. Bass, A.H. Ramos, G. Saksena, A.D. Cherniack, S.E. Schumacher, B. Tabak, S.L. Carter, N.H. Pho, H. Nguyen, R.C. Onofrio, A. Crenshaw, K. Ardlie, R. Beroukhim, W. Winckler, G. Getz, M. Meyerson, A. Protopopov, J. Zhang, A. Hadjipanayis, E. Lee, R. Xi, L. Yang, X. Ren, H. Zhang, N. Sathiamoorthy, S. Shukla, P.-C. Chen, P. Haseley, Y. Xiao, S. Lee, J. Seidman, L. Chin, P.J. Park, R. Kucherlapati, J. Todd Auman, K.A. Hoadley, Y. Du, M.D. Wilkerson, Y. Shi, C. Liquori, S. Meng, L. Li, Y.J. Turman, M.D. Topal, D. Tan, S. Waring, E. Buda, J. Walsh, C.D. Jones, P.A. Mieczkowski, D. Singh, J. Wu, A. Gulabani, P. Dolina, T. Bodenheimer, A.P. Hoyle, J. V. Simons, M. Soloway, L.E. Mose, S.R. Jefferys, S. Balu, B.D. O’Connor, J.F. Prins, D.Y. Chiang, D. Neil Hayes, C.M. Perou, T. Hinoue, D.J. Weisenberger, D.T. Maglinte, F. Pan, B.P. Berman, D.J. Van Den Berg, H. Shen, T. Triche Jr, S.B. Baylin, P.W. Laird, G. Getz, M. Noble, D. Voet, G. Saksena, N. Gehlenborg, D. DiCara, J. Zhang, H. Zhang, C.-J. Wu, S. Yingchun Liu, S. Shukla, M.S. Lawrence, L. Zhou, A. Sivachenko, P. Lin, P. Stojanov, R. Jing, R.W. Park, M.-D. Nazaire, J. Robinson, H. Thorvaldsdottir, J. Mesirov, P.J. Park, L. Chin, V. Thorsson, S.M. Reynolds, B. Bernard, R. Kreisberg, J. Lin, L. Iype, R. Bressler, T. Erkkilä, M. Gundapuneni, Y. Liu, A. Norberg, T. Robinson, D. Yang, W. Zhang, I. Shmulevich, J.J. de Ronde, N. Schultz, E. Cerami, G. Ciriello, A.P. Goldberg, B. Gross, A. Jacobsen, J. Gao, B. Kaczkowski, R. Sinha, B. Arman Aksoy, Y. Antipin, B. Reva, R. Shen, B.S. Taylor, T.A. Chan, M. Ladanyi, C. Sander, R. Akbani, N. Zhang, B.M. Broom, T. Casasent, A. Unruh, C. Wakefield, S.R. Hamilton, R. Craig Cason, K.A. Baggerly, J.N. Weinstein, D. Haussler, C.C. Benz, J.M. Stuart, S.C. Benz, J. Zachary Sanborn, C.J. Vaske, J. Zhu, C. Szeto, G.K. Scott, C. Yau, S. Ng, T. Goldstein, K. Ellrott, E. Collisson, A.E. Cozen, D. Zerbino, C. Wilks, B. Craft, P. Spellman, R. Penny, T. Shelton, M. Hatfield, S. Morris, P. Yena, C. Shelton, M. Sherman, J. Paulauskis, J.M. Gastier-Foster, J. Bowen, N.C. Ramirez, A. Black, R. Pyatt, L. Wise, P. White, M. Bertagnolli, J. Brown, T.A. Chan, G.C. Chu, C. Czerwinski, F. Denstman, R. Dhir, A. Dörner, C.S. Fuchs, J.G. Guillem, M. Iacocca, H. Juhl, A. Kaufman, B. Kohl III, X. Van Le, M.C. Mariano, E.N. Medina, M. Meyers, G.M. Nash, P.B. Paty, N. Petrelli, B. Rabeno, W.G. Richards, D. Solit, P. Swanson, L. Temple, J.E. Tepper, R. Thorp, E. Vakiani, M.R. Weiser, J.E. Willis, G. Witkin, Z. Zeng, M.J. Zinner, C. Zornig, M.A. Jensen, R. Sfeir, A.B. Kahn, A.L. Chu, P. Kothiyal, Z. Wang, E.E. Snyder, J. Pontius, T.D. Pihl, B. Ayala, M. Backus, J. Walton, J. Whitmore, J. Baboud, D.L. Berton, M.C. Nicholls, D. Srinivasan, R. Raman, S. Girshik, P.A. Kigonya, S. Alonso, R.N. Sanbhadti, S.P. Barletta, J.M. Greene, D.A. Pot, K.R. Mills Shaw, L.A.L. Dillon, K. Buetow, T. Davidsen, J.A. Demchok, G. Eley, M. Ferguson, P. Fielding, C. Schaefer, M. Sheth, L. Yang, M.S. Guyer, B.A. Ozenberger, J.D. Palchik, J. Peterson, H.J. Sofia, E. Thomson., Comprehensive molecular characterization of human colon and rectal cancer., Nature. 487 (2012) 330–337.

[58] Cancer Genome Atlas Research Network, P.S. Hammerman, M.S. Lawrence, D. Voet, R. Jing, K. Cibulskis, A. Sivachenko, P. Stojanov, A. McKenna, E.S. Lander, S. Gabriel, G. Getz, C. Sougnez, M. Imielinski, E. Helman, B. Hernandez, N.H. Pho, M. Meyerson, A. Chu, H.-J.E. Chun, A.J. Mungall, E. Pleasance, A. Gordon Robertson, P. Sipahimalani, D. Stoll, M. Balasundaram, I. Birol, Y.S.N. Butterfield, E. Chuah, R.J.N. Coope, R. Corbett, N. Dhalla, R. Guin, A. He, C. Hirst, M. Hirst, R.A. Holt, D. Lee, H.I. Li, M. Mayo, R.A. Moore, K. Mungall, K. Ming Nip, A. Olshen, J.E. Schein, J.R. Slobodan, A. Tam, N. Thiessen, R. Varhol, T. Zeng, Y. Zhao, S.J.M. Jones, M.A. Marra, G. Saksena, A.D. Cherniack, S.E. Schumacher, B. Tabak, S.L. Carter, N.H. Pho, H. Nguyen, R.C. Onofrio, A. Crenshaw, K. Ardlie, R. Beroukhim, W. Winckler, P.S. Hammerman, G. Getz, M. Meyerson, A. Protopopov, J. Zhang, A. Hadjipanayis, S. Lee, R. Xi, L. Yang, X. Ren, H. Zhang, S. Shukla, P.-C. Chen, P. Haseley, E. Lee, L. Chin, P.J. Park, R. Kucherlapati, N.D. Socci, Y. Liang, N. Schultz, L. Borsu, A.E. Lash, A. Viale, C. Sander, M. Ladanyi, J. Todd Auman, K.A. Hoadley, M.D. Wilkerson, Y. Shi, C. Liquori, S. Meng, L. Li, Y.J. Turman, M.D. Topal, D. Tan, S. Waring, E. Buda, J. Walsh, C.D. Jones, P.A. Mieczkowski, D. Singh, J. Wu, A. Gulabani, P. Dolina, T. Bodenheimer, A.P. Hoyle, J. V. Simons, M.G. Soloway, L.E. Mose, S.R. Jefferys, S. Balu, B.D. O’Connor, J.F. Prins, J. Liu, D.Y. Chiang, D. Neil Hayes, C.M. Perou, L. Cope, L. Danilova, D.J. Weisenberger, D.T. Maglinte, F. Pan, D.J. Van Den Berg, T. Triche Jr, J.G. Herman, S.B. Baylin, P.W. Laird, G. Getz, M. Noble, D. Voet, G. Saksena, N. Gehlenborg, D. DiCara, J. Zhang, H. Zhang, C.-J. Wu, S. Yingchun Liu, M.S. Lawrence, L. Zou, A. Sivachenko, P. Lin, P. Stojanov, R. Jing, J. Cho, M.-D. Nazaire, J. Robinson, H. Thorvaldsdottir, J. Mesirov, P.J. Park, L. Chin, N. Schultz, R. Sinha, G. Ciriello, E. Cerami, B. Gross, A. Jacobsen, J. Gao, B. Arman Aksoy, N. Weinhold, R. Ramirez, B.S. Taylor, Y. Antipin, B. Reva, R. Shen, Q. Mo, V. Seshan, P.K. Paik, M. Ladanyi, C. Sander, R. Akbani, N. Zhang, B.M. Broom, T. Casasent, A. Unruh, C. Wakefield, R. Craig Cason, K.A. Baggerly, J.N. Weinstein, D. Haussler, C.C. Benz, J.M. Stuart, J. Zhu, C. Szeto, G.K. Scott, C. Yau, S. Ng, T. Goldstein, P. Waltman, A. Sokolov, K. Ellrott, E.A. Collisson, D. Zerbino, C. Wilks, S. Ma, B. Craft, M.D. Wilkerson, J. Todd Auman, K.A. Hoadley, Y. Du, C. Cabanski, V. Walter, D. Singh, J. Wu, A. Gulabani, T. Bodenheimer, A.P. Hoyle, J. V. Simons, M.G. Soloway, L.E. Mose, S.R. Jefferys, S. Balu, J.S. Marron, Y. Liu, K. Wang, J. Liu, J.F. Prins, D. Neil Hayes, C.M. Perou, C.J. Creighton, Y. Zhang, W.D. Travis, N. Rekhtman, J. Yi, M.C. Aubry, R. Cheney, S. Dacic, D. Flieder, W. Funkhouser, P. Illei, J. Myers, M.-S. Tsao, R. Penny, D. Mallery, T. Shelton, M. Hatfield, S. Morris, P. Yena, C. Shelton, M. Sherman, J. Paulauskis, M. Meyerson, S.B. Baylin, R. Govindan, R. Akbani, I. Azodo, D. Beer, R. Bose, L.A. Byers, D. Carbone, L.-W. Chang, D. Chiang, A. Chu, E. Chun, E. Collisson, L. Cope, C.J. Creighton, L. Danilova, L. Ding, G. Getz, P.S. Hammerman, D. Neil Hayes, B. Hernandez, J.G. Herman, J. Heymach, C. Ida, M. Imielinski, B. Johnson, I. Jurisica, J. Kaufman, F. Kosari, R. Kucherlapati, D. Kwiatkowski, M. Ladanyi, M.S. Lawrence, C.A. Maher, A. Mungall, S. Ng, W. Pao, M. Peifer, R. Penny, G. Robertson, V. Rusch, C. Sander, N. Schultz, R. Shen, J. Siegfried, R. Sinha, A. Sivachenko, C. Sougnez, D. Stoll, J. Stuart, R.K. Thomas, S. Tomaszek, M.-S. Tsao, W.D. Travis, C. Vaske, J.N. Weinstein, D. Weisenberger, D.A. Wigle, M.D. Wilkerson, C. Wilks, P. Yang, J. John Zhang, M.A. Jensen, R. Sfeir, A.B. Kahn, A.L. Chu, P. Kothiyal, Z. Wang, E.E. Snyder, J. Pontius, T.D. Pihl, B. Ayala, M. Backus, J. Walton, J. Baboud, D.L. Berton, M.C. Nicholls, D. Srinivasan, R. Raman, S. Girshik, P.A. Kigonya, S. Alonso, R.N. Sanbhadti, S.P. Barletta, J.M. Greene, D.A. Pot, M.-S. Tsao, B. Bandarchi-Chamkhaleh, J. Boyd, J. Weaver, D.A. Wigle, I.A. Azodo, S.C. Tomaszek, M. Christine Aubry, C.M. Ida, P. Yang, F. Kosari, M. V. Brock, K. Rogers, M. Rutledge, T. Brown, B. Lee, J. Shin, D. Trusty, R. Dhir, J.M. Siegfried, O. Potapova, K. V. Fedosenko, E. Nemirovich-Danchenko, V. Rusch, M. Zakowski, M. V. Iacocca, J. Brown, B. Rabeno, C. Czerwinski, N. Petrelli, Z. Fan, N. Todaro, J. Eckman, J. Myers, W. Kimryn Rathmell, L.B. Thorne, M. Huang, L. Boice, A. Hill, R. Penny, D. Mallery, E. Curley, C. Shelton, P. Yena, C. Morrison, C. Gaudioso, J.M.S. Bartlett, S. Kodeeswaran, B. Zanke, H. Sekhon, K. David, H. Juhl, X. Van Le, B. Kohl, R. Thorp, N. Viet Tien, N. Van Bang, H. Sussman, B. Duc Phu, R. Hajek, N. Phi Hung, K.Z. Khan, T. Muley, K.R. Mills Shaw, M. Sheth, L. Yang, K. Buetow, T. Davidsen, J.A. Demchok, G. Eley, M. Ferguson, L.A.L. Dillon, C. Schaefer, M.S. Guyer, B.A. Ozenberger, J.D. Palchik, J. Peterson, H.J. Sofia, E. Thomson, P.S. Hammerman, D. Neil Hayes, M.D. Wilkerson, N. Schultz, R. Bose, A. Chu, E.A. Collisson, L. Cope, C.J. Creighton, G. Getz, J.G. Herman, B.E. Johnson, R. Kucherlapati, M. Ladanyi, C.A. Maher, G. Robertson, C. Sander, R. Shen, R. Sinha, A. Sivachenko, R.K. Thomas, W.D. Travis, M.-S. Tsao, J.N. Weinstein, D.A. Wigle, S.B. Baylin, R. Govindan, M. Meyerson, Comprehensive genomic characterization of squamous cell lung cancers., Nature. 489 (2012) 519–525.

[59] S.A. Forbes, D. Beare, H. Boutselakis, S. Bamford, N. Bindal, J. Tate, C.G. Cole, S. Ward, E. Dawson, L. Ponting, R. Stefancsik, B. Harsha, C.Y. Kok, M. Jia, H. Jubb, Z. Sondka, S. Thompson, T. De, P.J. Campbell, COSMIC: somatic cancer genetics at high-resolution., Nucleic Acids Res. 45 (2017) D777–D783.

[60] C. Dong, P. Wei, X. Jian, R. Gibbs, E. Boerwinkle, K. Wang, X. Liu, Comparison and integration of deleteriousness prediction methods for nonsynonymous SNVs in whole exome sequencing studies., Hum. Mol. Genet. 24 (2015) 2125–2137.

[61] Y. Yin, K. Kundu, L.R. Pal, J. Moult, Ensemble variant interpretation methods to predict enzyme activity and assign pathogenicity in the CAGI4 NAGLU (Human N-acetyl-glucosaminidase) and UBE2I (Human SUMO-ligase) challenges., Hum. Mutat. 38 (2017) 1109–1122.

[62] L.G. Martelotto, C.K. Ng, M.R. De Filippo, Y. Zhang, S. Piscuoglio, R.S. Lim, R. Shen, L. Norton, J.S. Reis-Filho, B. Weigelt, Benchmarking mutation effect prediction algorithms using functionally validated cancer-related missense mutations., Genome Biol. 15 (2014) 484.

[63] F. Gnad, A. Baucom, K. Mukhyala, G. Manning, Z. Zhang, Assessment of computational methods for predicting the effects of missense mutations in human cancers., BMC Genomics. 14 Suppl 3 (2013) S7. doi:10.1186/1471-2164-14-S3-S7.

[64] A.S. Kondrashov, S. Sunyaev, F.A. Kondrashov, Dobzhansky-Muller incompatibilities in protein evolution., Proc. Natl. Acad. Sci. U. S. A. 99 (2002) 14878–14883.

[65] C.D. McFarland, K.S. Korolev, G. V Kryukov, S.R. Sunyaev, L.A. Mirny, Impact of deleterious passenger mutations on cancer progression., Proc. Natl. Acad. Sci. U. S. A. 110 (2013) 2910–5. doi:10.1073/pnas.1213968110.

[66] D.T. Jones, D. Cozzetto, DISOPRED3: precise disordered region predictions with annotated protein-binding activity., Bioinformatics. 31 (2015) 857–863.

[67] M. Pajkos, B. Mészáros, I. Simon, Z. Dosztányi, Is there a biological cost of protein disorder? Analysis of cancer-associated mutations., Mol. Biosyst. 8 (2012) 296–307.

[68] D. Frishman, P. Argos, Knowledge-based protein secondary structure assignment., Proteins Struct. Funct. Genet. 23 (1995) 566–579.

[69] F. Eisenhaber, P. Argos, Improved strategy in analytic surface calculation for molecular systems: Handling of singularities and computational efficiency., J. Comput. Chem. 14 (1993) 1272–1280.

[70] F. Eisenhaber, P. Lijnzaad, P. Argos, C. Sander, M. Scharf, The double cubic lattice method: Efficient approaches to numerical integration of surface area and volume and to dot surface contouring of molecular assemblies., J. Comput. Chem. 16 (1995) 273–284.

[71] P.F. Jonsson, P.A. Bates, Global topological features of cancer proteins in the human interactome., Bioinformatics. 22 (2006) 2291–2297.

[72] K.-I. Goh, M.E. Cusick, D. Valle, B. Childs, M. Vidal, A.-L. Barabási, The human disease network., Proc. Natl. Acad. Sci. U. S. A. 104 (2007) 8685–90. doi:10.1073/pnas.0701361104.

[73] G. Kar, A. Gursoy, O. Keskin, Human cancer protein-protein interaction network: A structural perspective., PLoS Comput. Biol. 5 (2009) e1000601.

[74] J. Liu, J.R. Faeder, C.J. Camacho, Toward a quantitative theory of intrinsically disordered proteins and their function., Proc. Natl. Acad. Sci. U. S. A. 106 (2009) 19819–19823.

[75] A. Fornili, A. Pandini, H.-C. Lu, F. Fraternali, Specialized Dynamical Properties of Promiscuous Residues Revealed by Simulated Conformational Ensembles., J. Chem. Theory Comput. 9 (2013) 5127–5147.

[76] C.J. Oldfield, J. Meng, J.Y. Yang, M.Q. Yang, V.N. Uversky, A.K. Dunker, Flexible nets: disorder and induced fit in the associations of p53 and 14-3-3 with their partners., BMC Genomics. 9 (2008) S1.

[77] H. Nishi, M. Tyagi, S. Teng, B.A. Shoemaker, K. Hashimoto, E. Alexov, S. Wuchty, A.R. Panchenko, Cancer missense mutations alter binding properties of proteins and their interaction networks., PLoS One. 8 (2013) e66273.

[78] P. Blume-Jensen, T. Hunter, Oncogenic kinase signalling., Nature. 411 (2001) 355–365.

[79] S.R. Lamandé, J.F. Bateman, W. Hutchison, R.J. McKinlay Gardner, S.P. Bower, E. Byrne, H.H. Dahl, Reduced collagen VI causes Bethlem myopathy: a heterozygous COL6A1 nonsense mutation results in mRNA decay and functional haploinsufficiency., Hum. Mol. Genet. 7 (1998) 981–989.

[80] D. Seeliger, B.L. de Groot, Protein thermostability calculations using alchemical free energy simulations., Biophys. J. 98 (2010) 2309–2316.

[81] Z. Shi, J. Sellers, J. Moult, Protein stability and in vivo concentration of missense mutations in phenylalanine hydroxylase., Proteins Struct. Funct. Bioinforma. 80 (2012) 61–70.

[82] J.J. Manfredi, The Mdm2-p53 relationship evolves: Mdm2 swings both ways as an oncogene and a tumor suppressor., Genes Dev. 24 (2010) 1580–1589.

[83] J.K. Pritchard, Are rare variants responsible for susceptibility to complex diseases?, Am. J. Hum. Genet. 69 (2001) 124–137.

[84] R.D. Kumar, A.C. Searleman, S.J. Swamidass, O.L. Griffith, R. Bose, Statistically identifying tumor suppressors and oncogenes from pan-cancer genome-sequencing data., Bioinformatics. 31 (2015) 3561–3568.

[85] S. Chun, J.C. Fay, Identification of deleterious mutations within three human genomes., Genome Res. 19 (2009) 1553–1561.

[86] Y. Choi, G.E. Sims, S. Murphy, J.R. Miller, A.P. Chan, Predicting the functional effect of amino acid substitutions and indels., PLoS One. 7 (2012) e46688.

[87] W.C. Wong, D. Kim, H. Carter, M. Diekhans, M.C. Ryan, R. Karchin, CHASM and SNVBox: toolkit for detecting biologically important single nucleotide mutations in cancer., Bioinformatics. 27 (2011) 2147–2148.

[88] X. Liu, X. Jian, E. Boerwinkle, dbNSFP v2.0: a database of human non-synonymous SNVs and their functional predictions and annotations., Hum. Mutat. 34 (2013) E2393–2402.

[89] A.A. Canutescu, A.A. Shelenkov, R.L. Dunbrack, A graph-theory algorithm for rapid protein side-chain prediction., Protein Sci. 12 (2003) 2001–2014.

[90] B. Rost, C. Sander, Conservation and prediction of solvent accessibility in protein families, Proteins Struct. Funct. Genet. 20 (1994) 216–226. doi:10.1002/prot.340200303.

